# Pre-neuronal origins of monoamine and acetylcholine synthesis and transport

**DOI:** 10.64898/2026.10.09.757917

**Authors:** Yiannis Pyrris, George Diallinas, Alexandros A. Pittis

**Affiliations:** Institute of Molecular Biology and Biotechnology (IMBB), Foundation for Research and Technology-Hellas, Heraklion, Crete, Greece; Department of Biology, National and Kapodistrian University of Athens, Athens, Greece

## Abstract

Neuronal architecture is highly modular and individual components can be found in non-neuronal cell types or even neuronless animals, suggesting some functions predate neurons. However, the evolutionary origins of neurotransmitter synthesis, transport and synaptic vesicle biogenesis outside animals remain obscure. To our knowledge, no previous work has systematically addressed the function of neuronal protein homologs outside animals. Our phylogenetic analysis indicates that orthologs of synaptic vesicle (SV) neurotransmitter transporters (SLC18) and synthesis enzymes are widespread in fungi, which, together with animals and sister taxa, form the clade Opisthokonta. Comparing them with their synaptic counterparts can shed light on the emergence of key functions at the animal-fungi split. Here, we combine extensive phylogenetic profiling and statistical analysis with the generation of functional data for these protein families in fungi. We show that monoamine synthesis, orthologous to that of neurons, is present in fungi and places its origin in the opisthokont common ancestor. We also annotate the orthologs of SLC18 transporters showing that they retain an exporter profile and reside predominantly in the endolysosomal system. Building on this observation, we couple deep-learning localization predictions and GO enrichment analysis to show that the phylogenetic context of SVs is lysosomal throughout the *Opisthokonta* tree, with implications for their phylogenetic origin. In addition, we monitor fungal gene expression in the presence of monoamines showing that they modulate secondary metabolism, hinting at their physiological significance in a neuronless clade. We propose that the machinery associated with neurotransmission originated before animals.

## INTRODUCTION

The emergence of multicellularity and cell type diversification are central to early metazoan evolution(*1*). More specifically, the appearance of neuronal cell types paved the way for complex responses to environmental changes and behavioral adaptations in animals(*2*, *3*). Despite its evolutionary and physiological importance, the origins of neuronal machinery components and their integration in a ‘*protoneuron*’ remain poorly understood. Research shows that neuronal architecture is highly modular, characterised by the co-option of distinct functional and structural components(*4*, *5*). Among these components are the toolkits for monoamine and acetylcholine neurotransmitter (NT) synthesis and transport. Monoamines like serotonin and dopamine have been shown to regulate multiple behaviors in vertebrates, such as feeding, social interactions, mood, activity levels and motor skills(*6–8*). Trace amines like tyramine, tryptamine and histamine have dominant roles as arthropod NTs, implicated in behavior and photoreception(*9*, *10*). These trace amines can also be found in vertebrate nervous systems exhibiting newly emerging roles(*11*, *12*). Acetylcholine on the other hand is the major transmitter of the vertebrate neuromuscular junction(*13*).

Most monoamine transmitters are synthesised in one or two steps. Trace amines come directly from decarboxylation of amino acids (by aromatic amino acid decarboxylases–AADCs). Serotonin and dopamine are produced in a two-step fashion by hydroxylation of amino acids and subsequent decarboxylation of the precursors 5-HTP and L-DOPA respectively (aromatic amino acid hydroxylases–AAAHs and AADCs)(*14*, *15*). Mammalian L-DOPA decarboxylase (DDC) has the most broad specificity, decarboxylating L-DOPA, 5-HTP and has been shown to bind tryptophan(*16–18*). The *Drosophila* DDC is specific for L-DOPA and 5-HTP(*19*). Tyrosine decarboxylases (TDC) have been lost in vertebrates and decarboxylate tyrosine to tyramine(*9*, *20*). Histidine decarboxylase (HDC) is highly specific for histidine(*21*). AAAHs are also specialized with a degree of substrate overlap. Phenylalanine hydroxylase (PH) hydroxylates phenylalanine and tryptophan to a lower extent(*22*). Tyrosine hydroxylase (TH) hydroxylates tyrosine to L-DOPA but also phenylalanine(*22*). Tryptophan hydroxylase produces 5-HTP from tryptophan and can hydroxylate phenylalanine with much lower affinity(*23*). Acetylcholine is synthesised from acetylation of choline (choline acetyltransferase–ChAT)(*24*). ChAT is a homolog of carnitine acetyltransferase–CrAT(*25*). Monoamine and acetylcholine transmitters are synthesised in the cytoplasm and loaded into synaptic vesicles by synaptic vesicle transporters of the SLC18 family. These are the SLC18 A clade vesicular monoamine transporters (VMATs) and vesicular acetylcholine transporters (VAchTs). The vesicular polyamine transporters (VPATs) form the SLC18B clade(*26*). For a simplified overview of the pathways see fig. 1A.

**Fig. 1.**
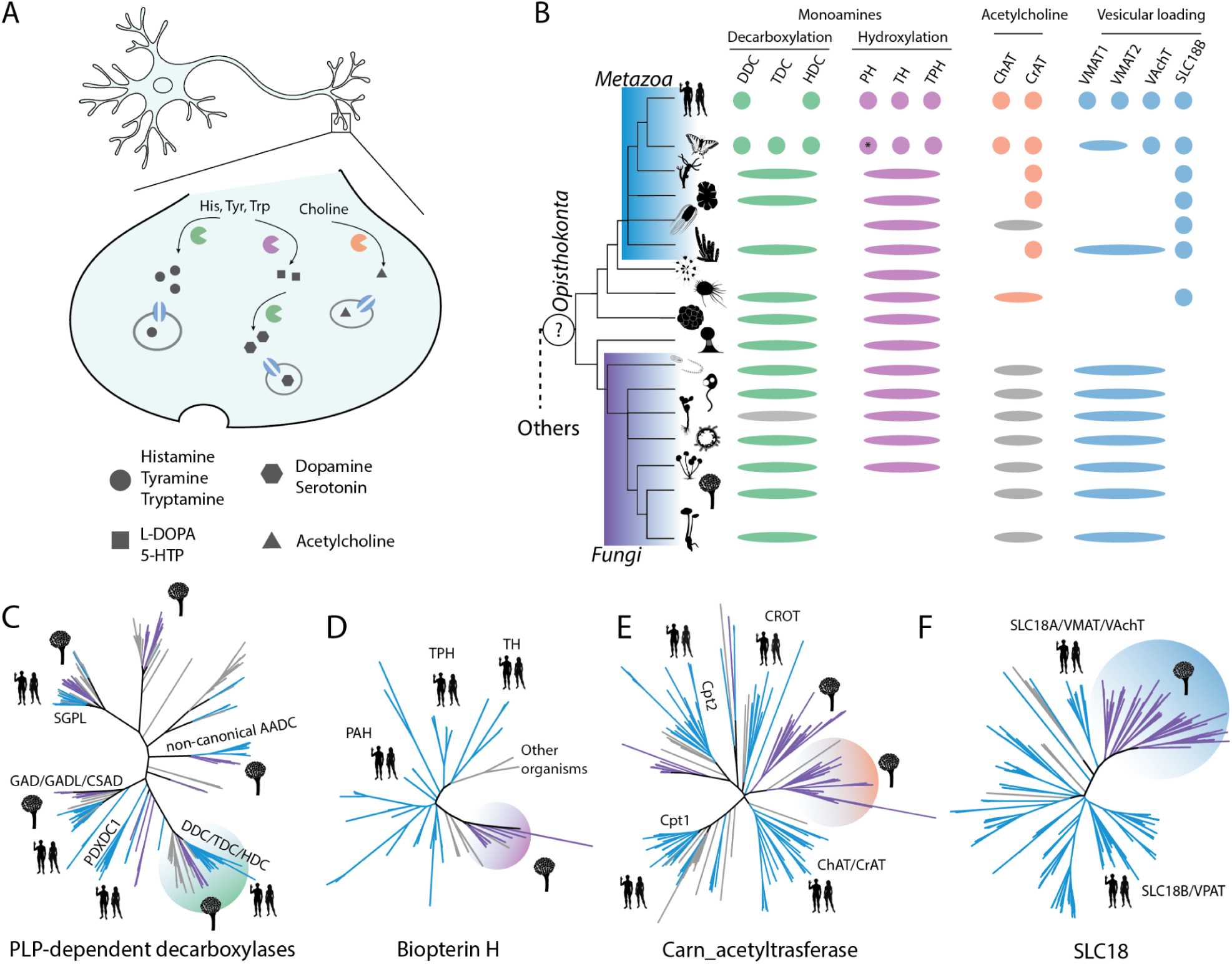
Homologs of the monoamine and acetylcholine toolkits are widespread. **A.** Simplified overview of monoamine and acetylcholine pathways in *bilaterian* neurons. **B.** Presence/absence matrix of monoamine and acetylcholine synthesis and transport genes. Elongated ellipse: ortholog to multiple proteins; gap: missing; grey ellipse: ortholog to both ChAT/CrAT and Cpt1. Visible taxonomic groups are from top to bottom: *Mammalia, Arthropoda, Cnidaria, Placozoa, Ctenophora, Porifera, Choanoflagellata, Filasterea, Ichthyosporea, Holomycota (Fonticula alba), Cryptomycota, Chytridiomycota, Blastocladiomycota, Zoopagomycota, Mucoromycota, Ascomycota, Basidiomycota.* All species can be seen in table S1. *: In *Drosophila* a phenylalanine hydroxylase is encoded by the *henna – hn* gene, which has ambiguous topology in our tree (fig. S3). **C.** Phylogeny of Pyridoxal_deC (PF00282) domain proteins, with the clade of the fungal AADC cloned highlighted. **D.** Phylogeny of Biopterin_H (PF00351) domain proteins, the fungal clade is highlighted. **E.** Phylogeny of Carn_acyltransf (PF00755) domain containing proteins, the position of the cloned gene is highlighted. **F.** Phylogeny of the SLC18 transporter clusters (see methods), with the fungal clade highlighted.

The evolutionary origins of the described NT-related machineries remain unclear. Importantly, they are present and have functions in animals without neurons. Yañez-Guerra *et al.*, working on *Placozoa*, simple neuronless animals with few cell types, showed that they respond behaviorally to the trace amine transmitters tyramine, tryptamine and phenylethylamine. They also demonstrated that placozoan GPCRs, related to *bilaterian* monoamine receptors, can be activated by trace amines(*27*). Work on sponges also identified behavioral responses to NTs(*28*, *29*). Zang *et al.* working on *Spongilla lacustris*, showed distinct responses to trace amines and modulation of GPCR signalling by them. They characterise a non-canonical monoamine synthesis and transport toolkit, consisting of aromatic amino acid decarboxylase homologs and Major Facilitator Superfamily MFS vesicular transporters(*30*). These studies challenge the idea that monoaminergic signalling and synthesis are *bilaterian* innovations and support a more ancient and promiscuous, in terms of machinery, origin of this system. To our knowledge the existence and function of the monoaminergic and cholinergic toolkits have not been systematically assessed outside animals. Thus, the question of how ancient some of their functions are remains largely unexplored.

Here we phylogenetically characterise the AADC, AAAH, ChAT/CrAT and SLC18 orthologs of animals, fungi and their unicellular relatives, constituting the clade *Opisthokonta*. We show that representatives for each protein family are present throughout *Opisthokonta*. These toolkits are extensively duplicated in *Bilateria*. Based on the most parsimonious evolutionary route, the common ancestor of animals and fungi had at least one representative of each family. To infer the functions in the common ancestor, we functionally characterise fungal AADC, ChAT/CrAT and SLC18 orthologs. We show that promiscuous trace amine synthesis is mediated by the fungal AADC. We propose that monoamine synthesis was present in the common ancestor of animals and fungi. This synthesis was mediated by the same machinery that *Bilateria* harbor in a duplicated and specialised form. We also demonstrate that fungal SLC18 proteins function as exporters (i.e. efflux transporters), which is in line with the directionality of synaptic vesicle transporters, and confer chemical resistance to strains overexpressing them. We show that they predominantly localise to the vacuoles/lysosomes and endosomes. We employ a large-scale phylogenomic approach, combining deep-learning localisation predictions and GO-term functional analysis for synaptic vesicle (SV) protein homologs across *Opisthokonta*. We propose that a large fraction of the SV proteome has a lysosomal origin, with contributions from the plasma membrane and to a lesser extent from other secretory pathway compartments, making up a hybrid organelle. By gathering expression data from *wild-type* strains grown in the presence of trace amines, we show that trace amines alter fungal gene expression. We observe modulation of fungal secondary metabolism and genes involved in biotic interactions. Collectively, our results support that elements of the cholinergic and monoaminergic systems are present in fungi and by inference in the common ancestor of *Opisthokonta*.

## RESULTS

### Phylogenies of monoamine and acetylcholine synthesis and transport homologs

We used a broad phylogenetic sampling focused on *Opisthokonta.* We included representatives from major metazoan lineages (*Bilateria*, *Cnidaria*, *Placozoa*, *Ctenophora* and *Porifera*), unicellular animal relatives, relatives of fungi and representatives from major fungal lineages (table S1). With this data, we generated a presence/absence matrix summarizing the phylogenetic profile of AADCs, AAAHs, ChAT/CrATs and SLC18 transporters (fig. 1B). To generate the matrix, we gathered all sequences corresponding to their characteristic Pfam profiles and reconstructed their phylogenies (Fig. 1C-F). For SLC18 transporters we included a clustering step, since their profile corresponds to the very large family of MFS transporters (methods). We named tree clades based on the presence of annotated *bilaterian* members.

The presence/absence profile shows a recurring duplication pattern for every family investigated. Proteins involved in monoamine and acetylcholine synthesis and transport have duplicated in the *bilaterian* lineage (Fig. 1B). This pattern suggests lineage specific adaptation and refinement of the system. Our results agree with a previous study that investigated the evolution of the monoaminergic system(*31*). However, we note that this view of protein family evolution provides a *bilaterian*-centric view. Monoaminergic and cholinergic system complexity appears higher in *bilaterian* animals, because their annotated proteins are used as a complexity measure. In our phylogenies we also observe multiple lineage specific duplications in other metazoan lineages, in the same or related protein families (fig. S2-S5). To our surprise, ctenophores, which lack an AADC ortholog, retain a non-canonical aromatic amino acid decarboxylase bearing the same domain as AADC, which has been previously studied in mushroom fungi (fig. S2 and discussion)(*32*, *33*). The analysis presented in this paper uses the *bilaterian* proteins as a reference.

Orthologs of the duplicated *bilaterian* proteins have a wide taxonomic distribution, with representatives being present in most *Opisthokonta* proteomes (fig. 1B). Interestingly, fungi possess at least one homolog for all families investigated here, while some lineages show independent expansion of these families (fig. S2-S5). Fungal AADCs and AAAHs are placed as a sister clade to the animal proteins (fig. 1C, D, fig. S2, S3). Duplications in *Bilateria* resulted in three different paralogs which are specialized for specific substrates. However, as noted in the introduction, these enzymes show a degree of functional overlap. This could suggest that they are derived from a promiscuous ancestor. Fungal ChAT/CrAT homologs are orthologous to both ChAT/CrAT and to Carnitine O-palmitoyltransferase 1–Cpt1 (fig. 1E). Notably, only *Bilateria* appear to have both copies of the ChAT/CrAT family with other *Metazoa* retaining only the CrAT enzyme. SLC18 transporters are expanded in *Bilateria*. Other *Metazoa*, except sponges which have SLC18A, retain only the SLC18B clade. Interestingly, Fungi appear to retain only the SLC18A clade of monoamine and acetylcholine transporters. Nevertheless, the exact clade topology could be different with fungal sequences placed as a sister to all animal SLC18 proteins. Note that fungi have multiple lineage specific expansions of this family (fig. S5). These results suggest that the last common ancestor of animals and fungi had homologs of the whole arsenal for monoamine and acetylcholine synthesis and transport. These families later expanded multiple times in animals.

### The fungal AADC can synthesise trace amines

The above data raised the question of which synthesis functions were present in the common ancestor and which evolved after the animal-fungi split. By comparing the functions of well annotated animal proteins with functions of their fungal homologs, information about their evolution can be gained. Functions that appear on both sides of the split were probably shared by the common ancestor, based on the assumption that the most probable evolutionary path is the one that requires the fewest steps (maximum parsimony).

To approach our question, we initially inspected the modelled fungal AADC Alphafold structure. The prediction exhibits a 3D-conformation that is highly similar to the human DDC and HDC crystal structures. Interestingly the binding site of the fungal protein shows high similarity to both human DDC and HDC (fig. 2A, B). To experimentally validate this observed similarity, we cloned and overexpressed the fungal AADC in the model fungus *Aspergillus nidulans*. The strain overexpressing the enzyme was grown under identical conditions to a *wild-type* strain and then both were subjected to targeted metabolomics, testing for neurotransmitter presence (fig. 2C, fig. S6). We found that AADC overexpression leads to increased tryptamine (9.2-fold, Hedges’ g = 24.3, Benjamini–Hochberg-adjusted q = 9.1 × 10⁻⁶), tyramine (6.1-fold, Hedges’ g = 3.4, q = 0.010) and histamine (4.5-fold, Hedges’ g = 2.2, q = 0.044) compared to the control (Welch’s t-test on log₁₀-transformed peak areas, FDR corrected across all compounds and comparisons). This result suggests that the enzyme can decarboxylate multiple amino acids and produce trace amines.

**Fig. 2.**
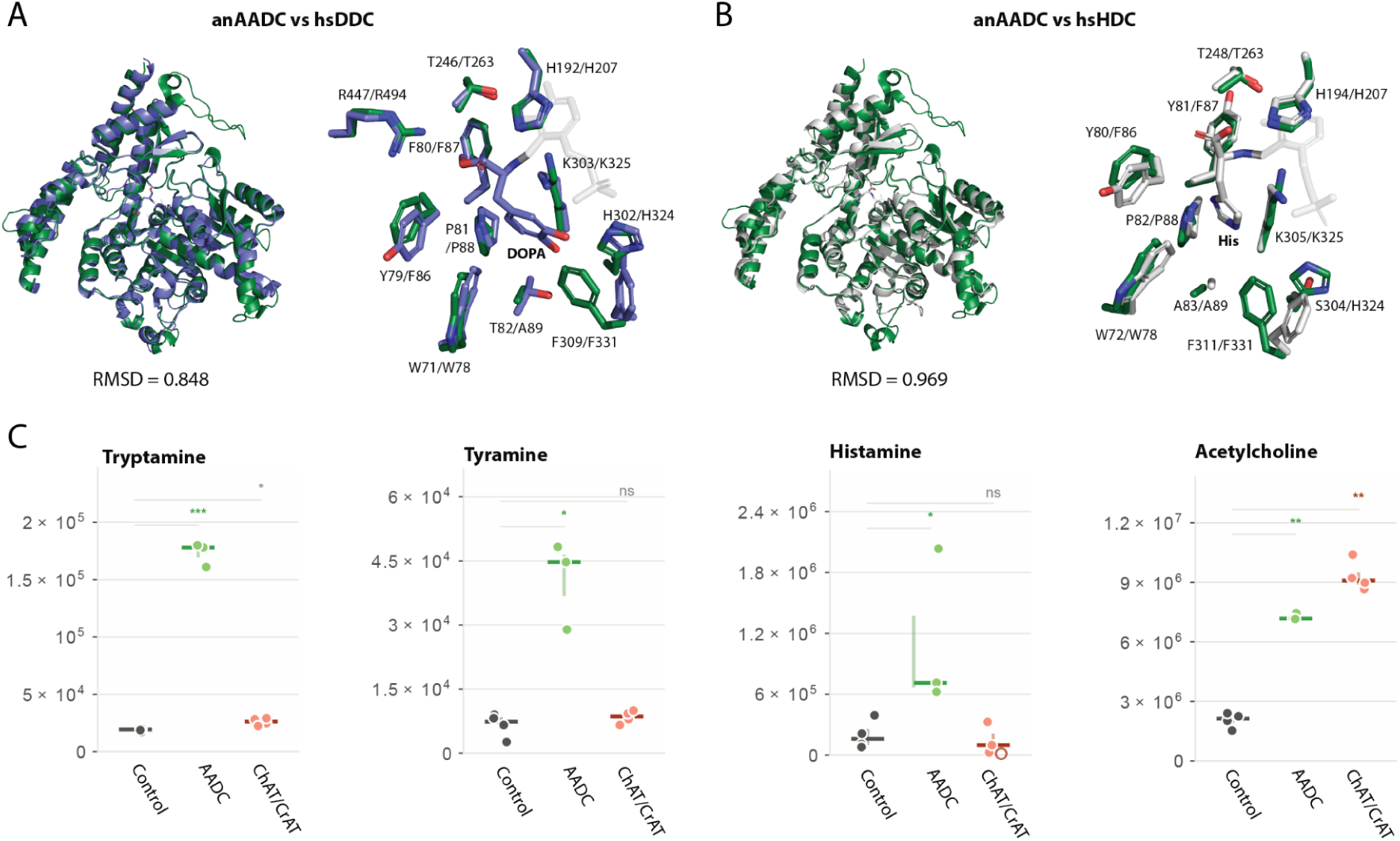
The fungal AADC can synthesise trace amines. **A.** Structure overlay and binding site comparison of the fungal AADC – green with the human DDC – blue (8ORA). **B.** with human HDC – grey (7EIW). All RMSD values are in Angstrom. L-DOPA methyl ester and histidine are part of the cited structures. The cofactor PLP can be seen with low-opacity grey. **C.** Relative metabolomic quantification of monoamines and acetylcholine. The y-axis is the relative peak area. Strains expressing the specified proteins are compared to a control strain. Stars = BH q of the Welch test on log₁₀ area versus the strain’s own control (*** q < 0.001, ** q < 0.01, * q < 0.05); coloured and bold = also at least 1.5-fold and above three times the QC coefficient of variation; grey = significant but below the effect-size or noise threshold.

The fungal AADC encompasses functions of the *bilaterian* HDCs and TDCs, while also being able to decarboxylate tryptophan like a DDC. Based on these results, promiscuous trace amine synthesis could be an ancestral trait of animals and fungi. Later duplication and specialisation events gave rise to the extant *bilaterian* enzymes, but homologous capacity for monoamine synthesis was there long before.

We also performed metabolomics on a strain overexpressing a fungal ChAT/CrAT homolog (fig. 2C). This strain did not exhibit elevated trace amine content, serving as another control for the specificity of our assay. However, it showed a 4.26-fold increase (Hedges’ g = 8.7, q = 0.001) in acetylcholine content, suggesting that the enzyme mediates acetylcholine synthesis. Note that the AADC overexpression strain also has increased acetylcholine content but to a lesser extent (3.36-fold, Hedges’ g = 6.91, q = 0.004). This suggests that protein overexpression could lead to an increase in intracellular acetylcholine and the magnitude of the result can partially be attributed to this.

### Fungal SLC18 transporter activity and localization

Fungal SLC18 proteins are nested within the SLC18 family, next to *bilaterian* monoamine and acetylcholine transporters (fig. 3A). Using our model system we overexpressed and GFP-tagged its native SLC18 proteins (five in total). Strains overexpressing the GFP-tagged transporters were subjected to growth tests on media containing the toxic compounds ethidium bromide and acriflavine, always compared to the control strain (fig. 3B). Overexpression of two of our transporters conferred chemical resistance to the respective strains. AN6387 (gene ID) overexpression led to transformants highly resistant in both treatments, while the AN7351 overexpression strain exhibited only acriflavine resistance (fig. 3C).

**Fig. 3.**
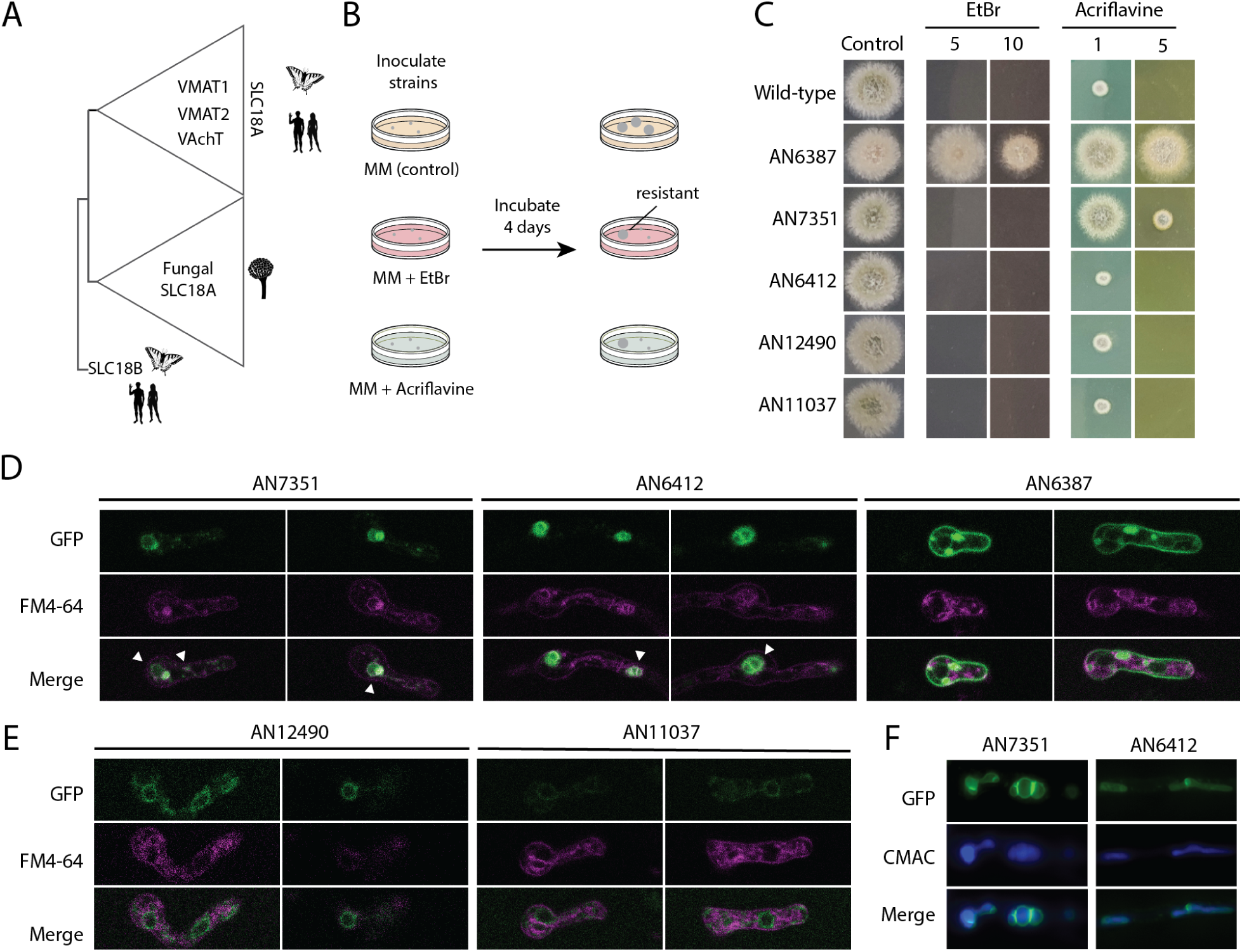
Fungal SLC18s confer chemical resistance and localise to the vacuole. **A.** Schematic depicting phylogenetic relations between fungal and *bilaterian* SLC18 proteins. **B.** Illustration of the growth assay testing for strain resistance to ethidium bromide (EtBr) and acriflavine. **C.** Growth tests of control strain (Wild-type) and strains overexpressing different GFP-tagged SLC18 proteins. The control condition is nitrate containing minimal medium (MM). Test conditions are MM containing 5ug/ml and 10ug/ml of EtBr and 1ug/ml and 5ug/ml of acriflavine. **D.** Confocal microscopy images of germlings overexpressing GFP-tagged transporters. Germlings were stained with FM4-64 which labels the endocytic pathway (endosomes and vacuoles). **E.** Same as D., but for ER-retained constructs. **F.** Colocalisation of the vacuolar transporters with the dye CMAC, which is internalised by vacuoles, visualised through epifluorescence microscopy.

Confocal microscopy revealed that AN7351 and AN6412 reside in the endolysosomal system of the fungus (lysosome/vacuole and endosomes), as they colocalize with the endocytic pathway dye FM4-64 staining the vacuolar membrane (fig. 3D). AN6387 had a different localisation profile mainly partitioning to the plasma membrane (PM) and characteristic perinuclear endoplasmic reticulum (ER) rings, that do not colocalise with FM4-64 (fig. 3D). Note the GFP signal present within the vacuoles and not on their membrane, suggesting degradation and not native targeting (fig. 3D). This has been shown for native versus degraded GFP-tagged vacuolar protein signals in plants(*34*). The last two transporters (AN12490 and AN11037) exhibit very low GFP signal and localise to the ER (fig. 3E). This is typical for misfolded membrane proteins, possibly due to the GFP-tag(*35*, *36*). The targeting profile of AN7351 and AN6412 was also validated using the dye CMAC, which accumulates inside the vacuole. The GFP signal decorates the membrane, while the dye localises inside the structure (fig. 3F).

Importantly, the rat VMAT2 (rVMAT2) also localises to the vacuolar membrane when heterologously expressed in yeast (fungal) cells, where it is able to confer resistance to ethidium bromide(*37*). A mammalian SV transporter has the propensity to traffic to the lysosome/vacuole when expressed in fungi. Two out of three successfully tagged fungal SLC18s also localize to the vacuole. This hints at a possible evolutionary link between SVs and lysosomes and a scenario where the ancestral SLC18 was a lysosomal/vacuolar resident protein, later incorporated into emerging SV-related organelles. Also, just like its fungal homologs, rVMAT2 mediates chemical resistance. Based on the localisation profiles of the resistance inducing transporters (AN6387 and AN7351) we propose that they function towards substrate export from the cell (AN6387) or import to the vacuole (AN7351), sequestering the chemicals. Moving molecules out of the cell is the same direction as importing them to the lumen of a secretory pathway organelle, like an SV or lysosome. That means that both *bilaterian* and fungal SLC18s function as exporters, consistent with the exporter profile being an ancestral trait of this transporter family.

### Lysosomes contributed to the emerging SV proteome

The possible phylogenetic implications of the SLC18 results prompted us to further investigate the origins of the SV proteome. We set out to predict the localisation of SV protein homologs at a large scale (Fig. 4A, methods). Briefly, we used a previously published dataset of rat resident synaptic vesicle proteins(*38*). Each protein in the dataset was used as a query for blastp searches against a database of proteomes representative of *Opisthokonta*. The hits, for all queries, for each organism were considered its SV protein homologs. Using DeepLoc2.0 we predicted the localisations of proteins in all proteomes. To investigate possible membrane donors for the emerging SV, we focused our analysis to transmembrane proteomes, by filtering all data for transmembrane proteins (DeepLoc2.0 transmembrane score ≥ 0.8). For groups of organisms (e.g. fungi, *Metazoa*), we compared the predicted localisations of SV protein homologs to the localisations of their whole transmembrane proteomes (fig. 4B, C, fig. S7).

**Fig. 4.**
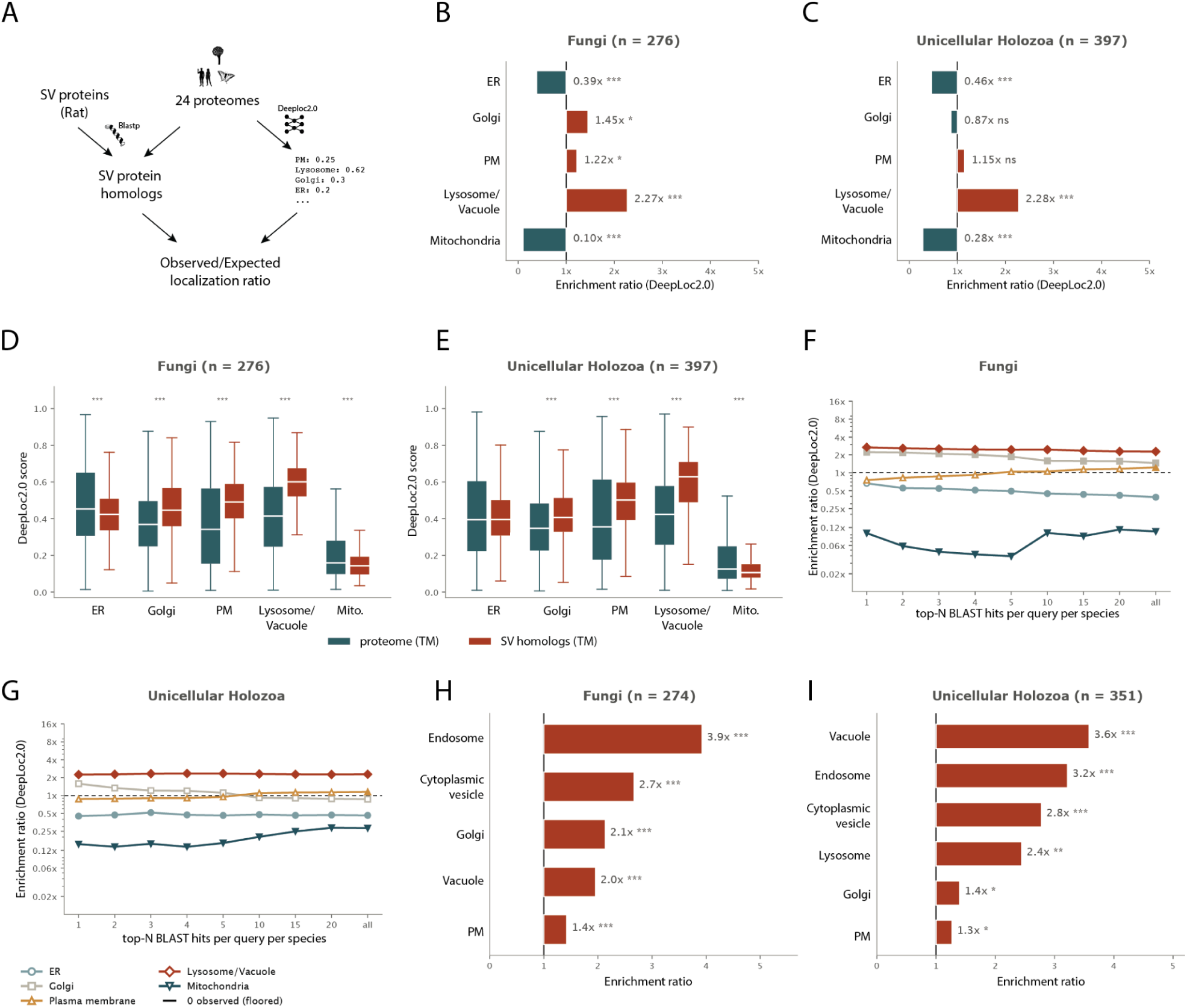
SV protein homologs have a major lysosomal component. **A.** Schematic of the pipeline used (for details see methods). **B, C.** Ratio of observed localisations to expected localisation for specific organelles for fungi (*A. nidulans, S. cerevisiae, B. dendrobatidis*) and unicellular holozoa (*S. rosetta, Monosiga brevicolis, C. owczarzaki, S. arctica*). **D, E.** Localisation probability distributions of SV homologs versus whole transmembrane proteomes. **F, G.** Ratio of observed localisations to expected localisation for specific organelles for fungi and unicellular holozoa in a sweep including different numbers of top hits. Empty plot points: no significant enrichment (methods). **H, I.** GO slim (cellular component) enrichment for SV homologs term counts versus expected term counts. Only proteins with GO annotation were used for calculation, that is why the n numbers differ from B, C. Stars denote adjusted q < 0.05 (*), < 0.01 (**) and < 0.001 (***).

We show that fungal SV protein homologs are >2 times enriched for predicted vacuolar/lysosomal localisation compared to their whole proteome background (n = 276; species-stratified hypergeometric resampling test, 20,000 draws, two-sided, Benjamini–Hochberg (BH)-adjusted q < 0.001). They also present more moderate enrichments for golgi (BH q = 0.026) and plasma membrane – PM (BH q = 0.026). Endoplasmic reticulum – ER (BH q < 0.001) and mitochondrial (BH q < 0.001) localisations are depleted (fig. 4B). Similar results are observed for unicellular animal relatives (*Filasterea, Icthysporea, Choanoflagellata*), with the lysosomal enrichment remaining stable (n = 397; BH q < 0.001) but more moderate magnitudes for other effects (fig. 4C). The core result remains stable for all different taxonomic groups tested (fig. S7).

For a more unbiased approach, instead of comparing localisation term outputs, we directly compared the DeepLoc2.0 output localisation probabilities (fig. 4D, E, fig. S8). Again, we observed differences with lysosome/vacuole, PM and Golgi probabilities being significantly higher in SV protein homologs compared to whole transmembrane proteomes (species-stratified permutation test on the mean probability, 4,000 permutations, two-sided, BH-adjusted q < 0.001 for each). ER and mitochondria had lower probabilities than the background (fig. 4D). The situation is similar for unicellular animal relatives (lysosome/vacuole 0.59 versus 0.42, PM 0.49 versus 0.40, Golgi 0.42 versus 0.37, mitochondrion 0.14 versus 0.22, all BH q < 0.001; ER 0.41 versus 0.42, BH q = 0.15; same test and correction) (fig. 4E) and other taxonomic groups (lysosome/vacuole higher in all nine groups, BH q < 0.001) (fig. S8).

The above analysis has an important drawback. By pooling all SV homologs, heavily duplicated families (one query that hits multiple homologs) will tend to be overrepresented, and so their localisations could dominate. To address this, we did the same analysis by keeping a specified number of blast hits per query, sorted by e-value (fig. 4F, G, S9). This way, for a cut-off of one blastp hit, an SV protein with ten homologs would be represented by only one sequence, same as a query with only three. The results appear to survive this stress test, even with a slight upward trend for lysosome/vacuole in fungi (2.27-fold with all hits to 2.66-fold with the top hit) and golgi enrichments (fungi 1.45-fold to 2.22-fold) with increasing phylogenetic relatedness – better e-value (fig. 4F, G). However the enrichment for plasma membranes appears to have the opposite trend (fungi 1.22-fold with all hits to 0.75-fold with the top hit). The above analysis was also done for each species separately (fig. S10), supporting the same results. We also skipped the transmembrane filtering step (fig. S11-S14), with the core results remaining essentially the same, with the difference of the ER showing moderate enrichment.

To complement the analysis done using the DeepLoc2.0 predictions, we also did enrichment analysis for GO-slim cellular component terms of transmembrane SV homologs compared to whole transmembrane proteomes (fig. 4H, I, S15). We show that the terms Vacuole, Lysosome and Endosome corresponding to the endolysosomal system are among the top enriched terms. Golgi and PM also show more moderate enrichments, closely following the prediction results. Analysis skipping the transmembrane filtering step gives similar results (fig. S16, S17). Even when applying the same hit cut-offs as for the localisation we observe the same results (fig. S18). Notably even when filtering for only the top blastp hit of SV proteins in mammals (many of which are SV proteins themselves), we get the same enrichment for lysosome terms in the localisation predictions (fig. S9). This could be explained as the effect of two factors: 1) DeepLoc2.0 tends to predict SV proteins as Lysosome/Vacuole since it has no category related to SVs, meaning that SV proteins have sequence features that categorise them as lysosomal by the model and 2) many SV proteins are in fact lysosomal proteins in mammals that partition in both organelles. Examples of this are vacuolar ATPase subunits. However, the result remains robust, in most groups, when removing the top query contributors to lysosomal localisation, showing that this result is not driven by only a few proteins (fig. S19).

We conclude that the phylogenetic context of resident SV proteins is enriched for lysosomal/vacuolar proteins. This suggests that lysosomes contributed to a large extent to SV emergence during the course of evolution. These acidified organelles are not purely degradation hubs but have multiple roles in metabolite storage and can even fuse with the plasma membrane in certain cell types (see discussion). Moreover we showed that the fungal SLC18 transporters, also reside predominantly in the vacuole. Importantly, contributions from the PM and other secretory pathway structures, especially Golgi, shaped the SV proteome. Thus, SVs could have emerged as a hybrid organelle.

### Trace amine presence modulates fungal secondary metabolism

We showed that trace amines are synthesised in *Aspergillus* via a native AADC homologous to the *bilaterian* enzymes. They have been shown to elicit behavioural responses in neuronless animals and induce signalling pathways (introduction). However fungi do not exhibit behaviours or motility like animals do. So we asked what is the physiological role of the native monoamines (histamine, tryptamine and tyramine) in fungi. To approach this question we collected bulk RNA expression data from cultures grown in minimal media (MM) containing varying concentrations of these amines, along with control cultures grown in MM (fig. 5A). We monitored gene expression changes compared to the control. GO enrichment analysis of upregulated differentially expressed (DE) genes revealed that trace amines even at the lowest concentration of 10uM elicit activation of genes associated with secondary metabolism terms (fig. 5B). To further investigate this result we predicted secondary biosynthetic clusters in the *Aspergillus nidulans* genome using antiSMASH. We found that both up- and downregulated DE genes tend to reside in predicted secondary biosynthetic clusters more than expected by chance (fig. 5C). We found clear activation of biosynthesis genes for orsellinic acid and its derivatives F-9775A/B, the antibiotic DHMBA and related compounds, and penicillin (fig. 5D). Importantly, several of these molecules are produced during co-culture of fungi with bacteria(*39*). Orsellinic acid is termed as an archetypal polyketide that is used as a substrate for the biosynthesis of other secondary metabolites(*39*). The same polyketide synthase that produces orsellinic acid (OrsA), also synthesises lecanoric acid(*39*, *40*), which has antibiotic and antifungal properties(*41*, *42*). Orsellinic acid downstream metabolites F-9775A/B are molecules that have a proposed role in interspecies chemical communication with bacteria(*43*). Penicillin is the first widely used antibiotic(*44*). DHMBA has been shown to mediate antagonistic interactions of fungi with gram-positive bacteria(*45*). It also can be used as a precursor for other antibiotics(*45*).

**Fig. 5.**
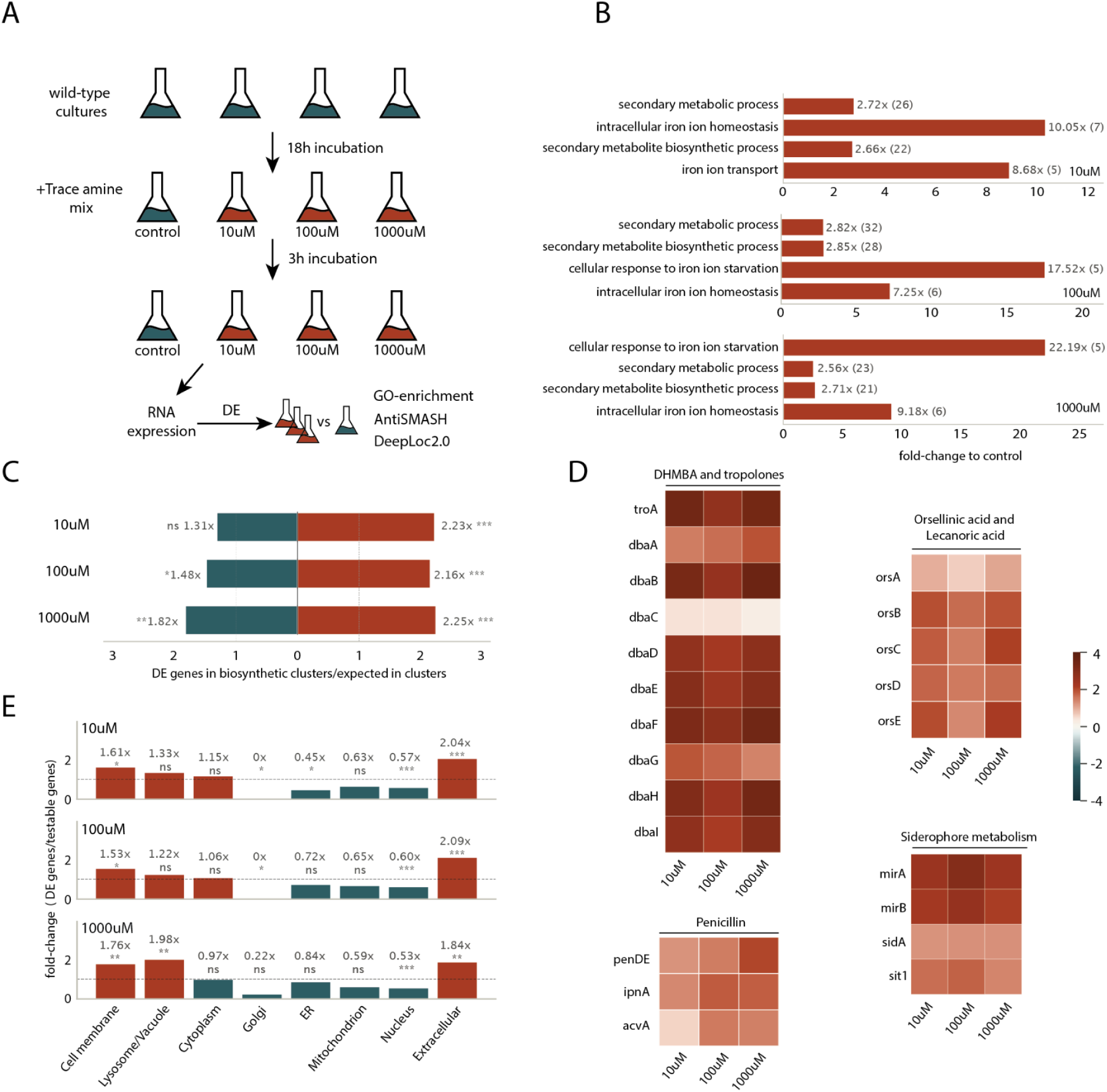
Trace amine presence induces secondary metabolism clusters. **A.** Schematic of the expression assay. **B.** GO enrichment (biological process) of trace amine exposed strains versus the control condition. **C.** Ratio of DE proteins residing in biosynthetic gene clusters versus the expected from the testable genome. **D.** Heatmaps of clusters or genes activated in trace amine presence. **E.** Enrichment of predicted protein localisations among DE genes relative to all testable genes. Stars denote adjusted q < 0.05 (*), < 0.01 (**) and < 0.001 (***).

Importantly in all conditions the upregulation of the DHMBA cluster coincides with the upregulation of the troA gene. This gene is involved in the production of tropolones from DHMBA(*45*). Tropolones are antibiotics with iron chelating properties. They have been shown to impede bacterial growth at a high concentration, but at low concentration they can be used by bacteria for iron uptake, making the interaction mutualistic(*45*). Relating to tropolones, we noticed that terms associated with iron homeostasis are also among upregulated DE genes (fig. 5B). Genes that have these terms are implicated in siderophore transport (*mirA*, *mirB*, *sit1*) and biosynthesis (*sidA*) (fig. 5D). These genes are again associated with mutualistic organism interactions (see discussion). Prediction of upregulated DE gene localisations, revealed that they are enriched for secreted/extracellular proteins compared to background genes for each condition (fig. 5E). This is also true for downregulated DE genes (fig. S20). These extracellular DE genes are enriched for carbohydrate metabolic functions (fig. S21).

The above results suggest that trace amines modulate fungal secondary metabolism. The most pronounced upregulated biosynthetic clusters are involved in antagonistic or symbiotic microbial interactions. Also many extracellular proteins are DE, suggesting altered interaction with the environment. Many bacteria have trace amine content(*46–48*). It is possible that *Aspergillus* translates trace amine presence as proximity to a microbial community. The validity of this claim requires further investigation. Whether these traits can be extended to the unicellular common ancestor of *Opisthokonta* or are lineage-specific adaptations remains unknown.

## DISCUSSION

Monoamines appear to have distinct signalling roles in animals whether they have neurons or not. Here we show that trace amines are also present in fungi and synthesised by machinery homologous to the one used in neurons. Additionally, we show that most homologs of the monoamine/acetylcholine synthesis and transport machinery are present throughout *Opisthokonta*, very often harboring their own lineage specific duplications. These duplications could suggest convergent evolution of these systems.

Zang *et al.* showed that the monoaminergic toolkit of sponges is non canonical, meaning that the families involved in synthesis and transport are not orthologs of the canonical AADCs and SLC18 transporters(*30*). They seem to be replaced by other PLP-dependent decarboxylases and SV2/SVOP/SLC17 transporters respectively(*30*). Here we show that, in our fungus, trace amines are synthesised by an ortholog of the *bilaterian* proteins, a canonical AADC, suggesting that the ancestor of animals and fungi exhibited canonical trace amine synthesis. Importantly, two other canonical fungal AADCs have been characterized. A *Basidiomycete* enzyme highly specific for 5-HTP and trp decarboxylation *in vitro* (*49*) and an enzyme from a *Penicillium* fungus with a broad specificity profile, capable of decarboxylating phe, tyr and trp but not histidine like the *A. nidulans* AADC(*50*). This suggests independent refinement of specificity in each lineage, similar to what is observed in animals. Together, these results hint that monoaminergic systems across the tree of life might be more flexible than previously thought. We imagine a scenario where monoamines could be derived from multiple pathways, exerting their signalling roles. Lineage specific duplication and adaptation, say, in sponges or *Bilateria*, later defined dominant synthesis and transport systems.

This idea is also corroborated by another observation. *Ctenophora* (comb jellies) have neurons but lack an AADC encoding gene (fig. 1B, fig. S2). However they harbor a non-canonical decarboxylase that is directly orthologous to a decarboxylase derived from *Psilocybe* mushrooms (fig. S2). This decarboxylase has been characterised in fungi and is capable of synthesising trace amines(*32*, *33*). Also ctenophores have an ortholog to the human Pdxdc1, which synthesises trace amines in sponges(*30*). If these proteins have the same function in comb jellies, they would provide another non-canonical pathway to monoamine synthesis. AAAH-encoding genes also have a wide distribution. However they appear lost in the lineage of mushrooms, molds and yeasts (*Dikarya*), which contains *Aspergillus* (fig. 1B, fig. S3). However they are present in early-diverging fungal lineages (fig. 1B), making the characterisation of these homologs interesting.

On the transport side, SLC18 transporters have their own lineage-specific expansions in fungi. Our system has five of them, with different localisation profiles (fig. 3D,E). This suggests selection for different functions. We found that two of them can function as resistance-conferring exporters when overexpressed. However we did not exhaustively characterise their specificity profiles, while the question of whether they are able to transport monoamines, like their *bilaterian* relatives, remains unresolved. Extensive characterisation of their transport capacities would shed more light to the evolutionary history of monoamine loading in SVs. However, as shown in the results section, they share common transport directionality and localisation features with the mammalian SLC18s. Like the rat VMAT2 they mostly partition to the lysosomes/vacuoles in fungi. This suggests that the ancestral SLC18 was a protein of the endolysosomal system.

Our large-scale approach to the origins of the synaptic vesicle proteome revealed that many homologs of SV proteins are lysosomal/vacuolar across the *Opisthokonta* tree (fig. 4). Related to our findings about the SV proteome and fungal SLC18s, SV precursor vesicles have been shown to colocalise with lysosomal markers during axonal transport in neurons(*51*). The same study also shows that lysosomes and SVs share the same kinesin adaptor for transport on microtubules (*51*). This suggests a mechanistic/biogenetic link between SVs and lysosomes that could stem from a common evolutionary origin. However, note that the above results are at variance with another study that shows no significant colocalisation between lysosomal and SV markers during biogenesis(*52*). Interestingly lysosomes have been shown to undergo regulated exocytosis in certain cell types. This exocytosis depends on calcium levels and is regulated by Syt7, a homolog of Syt1/2 which synaptic vesicles depend on(*53*, *54*). Syt7, which is abundant in lysosomes, also partitions in synaptic vesicles, where it has a role in regulating asynchronous release(*55–57*). Others have suggested that the SV pools within neurons should be seen as a continuum with their endolysosomal systems, as they overlap at multiple points, like SV protein endocytosis(*58*). Knocking out or overexpressing a Rab protein important for early endosome dynamics, has also been shown to alter SV pools in neurons(*59*, *60*).

The above evidence, combined with our result about the enrichment of lysosomal proteins among SV protein homologs, supports that the link between SVs and lysosomes is not only functional, but a result of common evolutionary origin. We propose that SVs and lysosomes are homologous structures. Our analysis also revealed substantial contributions from other compartments such as the Golgi apparatus and the PM, suggesting a role of multiple protein and membrane donors to their emergence. Importantly vacuoles are storage facilities for multiple secondary metabolites in plants and fungi(*61*, *62*). Plants also store monoamines and their derivatives in their vacuoles(*62*, *63*). These organelles could have functioned as an ancestral monoamine pool for the emerging SVs.

To try and understand the role of trace amines in fungal physiology, we monitored their effect on gene expression. Trace amines even at the lowest concentration led to a distinct modulation of fungal secondary metabolism, up- or downregulating multiple clusters. Notably we found multiple secreted proteins among the DE genes (fig. 5E) and the activation of clusters associated with antibiotic production (fig. 5D). As noted in the results section these antibiotics are produced when fungi are co-cultured with bacteria. Thus, in nature trace amine presence could hint to the composition of the microbial community, as they are found in multiple bacteria. Among the upregulated genes, we found siderophore biosynthesis enzymes and transporters. Siderophores are iron chelators that enhance iron uptake from the environment(*64*). Importantly, MirA and Sit1 are transporters able to utilize siderophores derived from bacteria, and not only the native ones(*65*, *66*). Based on the above we propose that trace amine presence can modulate biotic interactions between the fungus and other members of the microbial community.

We did not find any annotated clusters that produce secondary metabolites directly derived from any of the trace amines used. However native trace amine production is clearly present. Why does *Aspergillus* produce these compounds? Sensing them in the environment is distinct from their in-house production. Many of the modulated clusters we found are not annotated. It is possible that they could use trace amines as a substrate. Other fungi are able to produce indole alkaloids from trace amines such as tryptamine, although they mostly rely on alternative non-AADC decarboxylation routes(*67*, *68*). Tryptamine is also used for secondary metabolite synthesis in plants(*69*). Since AADCs are widespread in multiple fungal lineages (fig. S2) they could provide the substrates for secondary metabolite synthesis in multiple fungi.

The current study has some limitations. For one, ancestral state inference has an inherent degree of speculation. No one will ever be able to truly prove which functions were present in an extinct common ancestor. It is possible, although less probable, that trace amine synthesis by AADCs occurred independently in the fungal and animal lineages. Here we propose the most parsimonious route as the most probable. Also we do not stress our acetylcholine synthesis result from the ChAT/CrAT homolog, since the strain overexpressing AADC also has elevated acetylcholine content, suggesting that this could be an effect of protein overexpression induced stress. Lastly, our protein localisation analysis relies on predictions. Some of them will surely be wrong. To overcome this, we employed a conservative statistical analysis and multiple stress tests, across many lineages (fig. S8-S19).

Overall, our work shows that the machinery for monoamine/acetylcholine synthesis and transport is found in most opisthokont lineages. Specialisation via gene duplication appears to be the route followed by *Bilateria*. We show that trace amines are natively produced in fungi, by a canonical AADC, and produce a distinct transcriptional response. We also propose an evolutionary perspective on the origins of synaptic vesicles from the endolysosomal system. Thus many elements of the investigated systems may have been present before the split of animals and fungi.

## MATERIALS AND METHODS

### Proteomes used for phylogenetic analysis

The reference proteomes used for homology searches are in table S1.

### Sequence retrieval and clustering

The PFAM(*70*) profiles of interest: Pyridoxal_deC (PF00282), Biopterin_H (PF00351), Carn_acyltransf (PF00755) and MFS (PF07690), were used to collect proteins of interest. We used HMMER 3.3.2(*71*) hmmsearch with sequence and domain gathering thresholds set to the ones used by PFAM for each profile. Proteins containing the MFS domain (8,546 entries) were clustered using diamond v2.1.10.164(*72*) with --approx-id 20 and --member-cover 40. This resulted in 346 clusters. The two clusters containing known SLC18 proteins were used for downstream analysis.

### Alignment, trimming and phylogenetic tree reconstruction

Sequences were aligned with the MAFFT (v7.505) E-INS-I algorithm(*73*). Trimming was done using trimAl v1.5.rev0 with -gt 0.1(*74*). Phylogenetic trees were constructed using IQ-TREE version 3.0.1 with 1000 ultrafast bootstraps, and ModelFinder for model selection(*75*, *76*).

### Media, strains and growth conditions

Standard complete (CM) and minimal media (MM) for *A. nidulans* growth were used. Media and supplemented auxotrophies were used at the concentrations given in http://www.fgsc.net(77). Glucose 1% (w/v) was used as a carbon source. 10 mM sodium nitrate (NO_3_) was used as standard nitrogen source. All media and chemical reagents were obtained from Sigma-Aldrich (Life Science Chemilab SA, Hellas) or AppliChem (Bioline Scientific SA, Hellas). An *Aspergillus nidulans FSGC A4 pantoB100 bioA1* mutant strain was the recipient strain in transformations with plasmids carrying gene constructs based on complementation of the pantothenic acid auxotrophy *pantoB100*(*78*). *A. nidulans* protoplast isolation and transformation was performed as previously described(*79*). Growth tests were performed at 25°C for 96 h, at pH 6.8.

### Standard molecular biology manipulations and plasmid construction

Genomic DNA extraction from *A. nidulans* was performed as described in FGSC (http://www.fgsc.net). Plasmids, prepared in *E. coli*, and DNA restriction or PCR fragments were purified from agarose 1% gels with the Nucleospin Plasmid Kit or Nucleospin ExtractII kit, according to the manufacturer’s instructions (Macherey-Nagel, Lab Supplies Scientific SA, Hellas). Standard PCR reactions were performed using KAPATaq DNA polymerase (Kapa Biosystems). PCR products used for cloning, sequencing and re-introduction by transformation in *A. nidulans* were amplified by a high-fidelity KAPA HiFi HotStart Ready Mix (Kapa Biosystems) polymerase. DNA sequences were determined by VBC-Genomics (Vienna, Austria). The principal vector used for most *A. nidulans* mutants is a modified pGEM-T-easy vector carrying a version of the *gpdA* promoter, the *trpC* 3’ termination region and the *panB* selection marker. Gene IDs for AADC: AN10299; ChAT/CrAT: AN6279

### Fluorescence microscopy

Samples for standard epifluorescence microscopy were prepared as previously described(*80*). In brief, sterile 35 mm l-dishes with a glass bottom (Ibidi, Germany) containing liquid minimal media supplemented with NaNO_3_ and 1% glucose were inoculated from a spore solution and incubated for 16 h at 25°C. The images were obtained using a Leica SP8 inverted confocal microscope or an inverted Zeiss Axio Observer Z1 equipped with an Axio Cam HR R3 camera. Image processing was done using the Fiji (ImajeJ)(*81*) software while further processing of the TIFF files was made using Adobe Photoshop CS3 software for brightness adjustment, rotation, alignment and annotation. Brightness adjustments were made equally in all images. Labelling with FM4-64 was done modifying the protocol from (*82*). Briefly, strains of interest were incubated with the dye for 20 minutes (4°C), another 15 minutes at room temperature and then washed with liquid minimal medium, before observation. CMAC staining was done as in (*83*).

### Metabolomics

#### Wild-type Aspergillus nidulans

FSGCA4 strains and strains overexpressing genes of interest were inoculated in minimal media. Incubation followed at 25℃ for 18h in an orbital incubator (150 rpm). Mycelia were collected and snap-frozen in liquid nitrogen. Mortar and pestle were used to grind the mycelia to fine dust. Metabolite extraction and metabolomics were handled by the EMBL, Heidelberg metabolomics core facility. Each condition had ≥ 3 biological replicates. Appropriate minimal medium blanks were used to determine possible background in the medium. Statistical significance of differences compared to control was computed using Benjamini–Hochberg FDR-adjusted q-value (q < 0.05) taken from a Welch’s t-test on log₁₀(peak area) for each compound. Effect sizes are reported as Hedges’ g computed on log₁₀-transformed peak areas.

### Structure alignment and visualisation

The structure of the fungal AADC was downloaded from the Alphafold Structure Database(*84*, *85*). All structures were visualised using The PyMOL Molecular Graphics System, Version 2.5.0, Schrödinger, LLC. Structures were aligned using the PyMOL alignto command to calculate RMSD values.

### Query set and homolog identification for localisation prediction analysis

A previously published set of 119 (after removing sequences < 50aa that are not reliable for homology searches) resident synaptic vesicle (SV) proteins of *Rattus norvegicus* (*38*) was used as the query set. Each protein was searched with BLASTP(*86*) against a database of 24 opisthokont proteomes. The proteomes were grouped as Mammalia (*Homo sapiens*, *Mus musculus*, *R. norvegicus*), Arthropoda (*Drosophila melanogaster*), Cnidaria (*Nematostella vectensis*, *Hydra vulgaris*, *Paramuricea clavata*), Placozoa (*Trichoplax adhaerens*, *Trichoplax* sp. H2), Ctenophora (*Mnemiopsis leidyi*, *Bolinopsis microptera*, *Hormiphora californensis*), Porifera (*Amphimedon queenslandica*, *Sycon ciliatum*, *Corticium candelabrum*, *Halichondria panicea*), unicellular Holozoa (Choanoflagellata: *Monosiga brevicollis*, *Salpingoeca rosetta*; Filasterea: *Capsaspora owczarzaki*; Ichthyosporea: *Sphaeroforma arctica*), Fungi (*Aspergillus nidulans*, *Batrachochytrium dendrobatidis*, *Saccharomyces cerevisiae*) and unicellular Holomycota (*Fonticula alba*). Searches used an E-value threshold of 10⁻⁴ and retained up to 500 target sequences per query. Hits were collapsed to the best alignment per query–subject pair. Hits corresponding to the query protein itself were excluded as self-hits: the identical accession in the rat proteome (96 of 119 queries) and, for 23 queries, the same protein under a second rat accession (any rat hit with ≥ 99% sequence identity to the query). For each species, the distinct BLASTP subjects across all queries formed its set of SV protein homologs.

### Localization prediction and transmembrane filtering

Subcellular localization was predicted with DeepLoc 2.0(*87*) for every protein in all 24 proteomes. DeepLoc 2.0 is a multi-label classifier, so a protein was counted toward a compartment whenever that compartment was among its predicted labels. Proteins with a DeepLoc 2.0 transmembrane score ≥ 0.8 were classified as transmembrane. In the main analysis, both SV homologs and background were restricted to transmembrane proteins, so that the homologs of each species were compared with that species’ own transmembrane proteome. The analysis was repeated without this filter, against whole proteomes. In both cases the background included the homologs.

### Enrichment of predicted localizations

Each taxonomic group was analyzed separately, with a weighted expectation to account for differences in each species’ contribution to the homologs. For each organism group (like fungi) and each compartment the number of SV protein homologs localising to the compartment was summed for all organisms within the group. This is the observed count. The expected count for each organism is the fraction of proteins predicted to localise to that compartment in the whole proteome, multiplied by the number of homologs this organism contributed. The sum of these counts is the expected count of the group. The fold-change is the ratio of the observed to the expected count. For transmembrane proteome analysis the transmembrane filter (see methods) was applied equally to the background and to the homologs. Significance was assessed against a species-stratified (for each species the number of proteins contributed as homologs to the group was drawn) null in which proteins were drawn without replacement from the background of each species and pooled. This null is a sum of hypergeometric variables, and it was sampled 20,000 times. Two-sided *P* values were computed as twice the smaller empirical tail, (1 + *b*)/(*B* + 1), capped at 1. B = 20,000 draws, and b = the number of draws at least as extreme as the observed count, counted separately for each tail. They were corrected across compartments by the Benjamini–Hochberg procedure. The same analysis was run for each species individually.

### Comparison of localization probabilities/scores

To avoid thresholding the predictions, the mean DeepLoc 2.0 probability for each compartment was compared between SV homologs and the transmembrane background. Significance was assessed with the same species-stratified design, using 4,000 random draws of background proteins per species. Two-sided *P* values were corrected across compartments by the Benjamini–Hochberg procedure.

### Number of BLASTP hits per query

Large gene families can contribute many homologs per query and dominate a pooled homolog set. To control for this, the enrichment analysis was repeated retaining only the top *N* BLASTP hits per query per species (*N* = 1, 2, 3, 4, 5, 10, 15, 20 or all), ranked by E-value with bit score breaking ties. Hits were ranked after self-hits had been removed. At *N* = 1, each query contributes at most one homolog per species, regardless of family size.

### Query jackknife

To test whether the lysosome/vacuole enrichment depended on a few query proteins, queries were ranked by the number of transmembrane, lysosome/vacuole-predicted homologs they contributed in each group. They were then removed cumulatively, up to 20 queries, and the enrichment ratio and one-sided *P* value (5,000 draws) were recomputed after each removal.

### GO-slim enrichment analysis

Gene Ontology (GO) annotations were obtained from UniProtKB(*88*) through the UniProt REST API, including all evidence codes. Proteome accessions were mapped to UniProt entries directly or through RefSeq cross-references. Eight proteomes had no usable GO annotation: *M. leidyi*, *B. microptera*, *H. californensis*, *S. ciliatum*, *C. candelabrum*, *H. panicea*, *N. vectensis* and *Trichoplax* sp. H2. The GO-slim analysis was therefore performed on the remaining 16 proteomes, with Ctenophora absent and Porifera represented by *A. queenslandica* alone. Annotations were propagated to all ancestor terms through *is_a* and *part_of* relations of the GO release of 26 July 2026 (go-basic)(*89*). They were then mapped to the cellular-component terms of the goslim_generic subset.

Only proteins carrying at least one cellular-component annotation, homologs and background alike, were included. Proteins without such annotation cannot carry any of the terms being tested. Enrichment ratios were computed with the same species-weighted expectation, against the transmembrane background and against the whole annotated proteome. Terms covering fewer than 10 background proteins, or more than 95% of the background, were not tested. The stratified hypergeometric null distribution was computed exactly by convolving the per-species hypergeometric distributions. Two-sided *P* values were corrected across the tested terms of each group by the Benjamini–Hochberg procedure. The analysis was repeated over the same range of BLASTP hits per query.

### Culture in trace amine presence and RNA extraction

A *wild-type Aspergillus nidulans* FSGCA4 strain was inoculated in minimal media. Incubation followed at 25°C for 18h in an orbital incubator (150 rpm). After mycelium was formed (18h), 0uM, 10uM, 100uM and 1000uM final concentrations of a trace amine mix (histamine, tyramine, tryptamine) were added to the flasks. That resulted in 4 conditions total as shown in fig.5A. Each condition had 4 biological replicates. After 3h incubation in the same conditions, mycelia were collected and snap-frozen in liquid nitrogen. Mortar and pestle were used to grind the mycelia to fine dust. RNA extraction followed using the ZymoResearch *Quick*-RNA Microprep Kit (Cat: R1050). Quality control, library preparation and RNA sequencing were done by BMKgene (https://www.bmkgene.com/).

### RNA sequencing read processing, alignment and quantification

Adapters and low-quality bases were removed with Trim Galore (version 2.2.0)(*90*) in paired-end mode, with a Phred quality cutoff of 20, a minimum read length of 20 nt and automatic adapter detection. Trimmed reads (20.2 to 26.3 million pairs per sample) were aligned to the *A. nidulans* FGSC A4 genome (assembly ASM1142v1, Ensembl Fungi release 62) with STAR 2.7.11b(*91*), using the Ensembl gene annotation for splice junctions. Between 91.3% and 97.8% of read pairs mapped uniquely. Read pairs were assigned to the 10,828 annotated genes with featureCounts (Subread 2.1.1)(*92*), counting fragments rather than individual mates (-p --countReadPairs) in unstranded mode (-s 0).

### Differential expression

Differential expression was tested with PyDESeq2 0.5.4(*93*). A single model with the design ∼ condition was fitted to the raw counts of all 16 samples, with the control as the reference level. Each trace-amine concentration was compared with the control by a Wald test. *P* values were adjusted by the Benjamini–Hochberg procedure(*94*) after the default independent filtering and Cook’s-distance outlier filtering of DESeq2. Genes with an adjusted *P* value were called testable (9,223, 9,429 and 9,636 genes at 10 uM, 100 uM and 1000 uM). A gene was called differentially expressed (DE) when its adjusted *P* was below 0.05 and its |log2 fold change| (maximum-likelihood estimate, without shrinkage) was 1 or more. Up- and down-regulated genes were analyzed separately in the enrichment tests below, except where stated.

### Gene annotation

Gene identifiers (ANIA_xxxxx) were linked to UniProtKB accessions by combining the gene-name cross-references of the UniProt *A. nidulans* FGSC A4 entries with the synonym field of the UniProt-GOA annotation file for taxon 227321 (generated 10 April 2026). This mapped 10,517 of 10,828 genes (97.1%). GO terms were taken from the same annotation file, with all evidence codes, and the ontology from go-basic.

### GO enrichment analysis

GO over-representation was tested with GOATOOLS 1.6.5(*95*), separately for the biological process, molecular function and cellular component ontologies and separately for up- and down-regulated genes at each concentration. Gene annotations were propagated to all ancestor terms through *is_a* relations. For each term, the number of DE genes annotated to it was compared with the number expected from the population by Fisher’s exact test (two-sided), and *P* values were corrected by the Benjamini–Hochberg procedure across all terms tested within each ontology. Terms with a corrected *P* below 0.05 and more DE genes than expected were reported as enriched.

### Prediction of secondary-metabolite biosynthetic gene clusters

Biosynthetic gene clusters (BGCs) were predicted with antiSMASH 8.0.4(*96*) in fungal mode (--taxon fungi --minimal) on the ASM1142v1 genome, using the Ensembl gene models converted to GFF3 with gffread(*97*). This identified 61 regions containing 1,292 genes.

### Enrichment of DE genes in biosynthetic gene clusters

For each concentration and direction, we tested whether DE genes lay inside predicted BGCs more often than expected by chance. The background was the testable genes of that comparison (*N*), of which *K* lay inside a BGC. If *k* of the *n* DE genes lay inside a BGC, the fold enrichment was (*k*/*n*)/(*K*/*N*). Significance was assessed against a null in which *n* genes were drawn without replacement from the *N* testable genes, sampled 10,000 times. The one-sided *P* value was (1 + *b*)/(*B* + 1), where *b* is the number of draws with at least *k* BGC genes and *B* = 10,000. The exact hypergeometric *P* value was computed alongside and agreed with every call. *P* values were corrected by the Benjamini–Hochberg procedure across the tests of the full experiment. For individual clusters, the log2 fold change of every member gene was plotted.

### Enrichment of predicted localizations among DE genes

For each concentration, direction and compartment *c*, the enrichment ratio was the fraction of DE genes assigned to *c* divided by the fraction of testable genes assigned to *c*. Significance was assessed against a null in which the same number of genes as the DE set was drawn without replacement from the testable genes, sampled 10,000 times (fixed random seed). Two-sided *P* values were computed as twice the smaller empirical tail, 2 × (1 + *b*)/(*B* + 1), where *b* is the number of draws with a ratio at least as extreme in that tail, capped at 1. *P* values were corrected by the Benjamini–Hochberg procedure across all condition × compartment tests in each figure.

### GO enrichment of extracellular DE proteins

To describe the extracellular DE proteins, DE genes (up- and down-regulated together) whose proteins were predicted to be extracellular were tested for GO over-representation. The test was the same as above, except that the population was the testable genes of each comparison.

## Author contributions

Conceptualization: Y.P., A.A.P.

Methodology: Y.P.

Investigation: Y.P.

Visualization: Y.P.

Supervision: A.A.P., G.D.

Writing—original draft: Y.P.

Writing—review & editing: Y.P., A.A.P., G.D.

We acknowledge the support of the EMBL Metabolomics Core Facility (MCF) and especially Bernhard Drotleff in the acquisition and analysis of liquid chromatography-mass spectrometry data.

