## Supplementary material for "Pre-neuronal origins of monoamine and acetylcholine synthesis and transport"

Yiannis Pyrris *et al.*

**This PDF file includes:**

Figs. S1 to S21

**Table S1.**

**Reference proteomes used for phylogenetic analysis.** Species, taxonomic group, NCBI taxon ID and UniProt reference proteome identifier for all proteomes used in the homology searches.

| Name | Group | Taxon ID | Reference proteome |
|---|---|---|---|
| *Aspergillus nidulans* FGSC A4 | Ascomycota | 227321 | UP000000560 |
| *Aspergillus fumigatus* Af293 | Ascomycota | 330879 | UP000002530 |
| *Ophiocordyceps camponoti-floridani* | Ascomycota | 2030778 | UP000562929 |
| *Saccharomyces cerevisiae* S288C | Ascomycota | 559292 | UP000002311 |
| *Akanthomyces lecanii* RCEF 1005 | Ascomycota | 1081108 | UP000076881 |
| *Gonapodya prolifera* JEL478 | Chytridiomycota | 1344416 | UP000070544 |
| *Schizosaccharomyces pombe* 972h- | Ascomycota | 284812 | UP000002485 |
| *Amanita muscaria* Koide BX008 | Basidiomycota | 946122 | UP000054549 |
| *Agaricus bisporus* | Basidiomycota | 597362 | UP000008493 |
| *Wolfiporia cocos* | Basidiomycota | 742152 | UP000218811 |
| *Postia placenta* | Basidiomycota | 670580 | UP000194127 |
| *Pleurotus ostreatus* PC9 | Basidiomycota | 5322 | UP000623687 |
| *Psilocybe cubensis* | Basidiomycota | 181762 | UP000664032 |
| *Mucor circinelloides* 1006PhL | Mucoromycota | 1220926 | UP000014254 |
| *Rhizophagus clarus* HR1 | Mucoromycota | 94130 | UP000247702 |
| *Rhizopus microsporus* ATCC 52813 | Mucoromycota | 1340429 | UP000242254 |
| *Allomyces macrogynus* ATCC 38327 | Blastocladiomycota | 578462 | UP000054350 |
| *Rozella allomycis* CSF55 | Cryptomycota | 988480 | UP000030755 |
| *Basidiobolus meristosporus* CBS 931.73 | Zoopagomycota | 1314790 | UP000193498 |
| *Chytriomyces confervae* CBS 675.73 | Chytridiomycota | 246404 | UP000320333 |
| *Batrachochytrium dendrobatidis* JAM81 | Chytridiomycota | 684364 | UP000007241 |
| *Fonticula alba* | Holomycota | 691883 | UP000030693 |
| *Monosiga brevicollis* | Holozoa | 81824 | UP000001357 |
| *Salpingoeca rosetta* ATCC 50818 / BSB-021 | Holozoa | 946362 | UP000007799 |
| *Sphaeroforma arctica* JP610 | Holozoa | 667725 | UP000054560 |
| *Capsaspora owczarzaki* ATCC 30864 | Holozoa | 595528 | UP000008743 |
| *Bolinopsis microptera* | Ctenophora | 2820187 | Schultz et al. (2023) |
| *Mnemiopsis leidyi* | Ctenophora | 27923 | UP001729289 |
| *Amphimedon queenslandica* | Porifera | 400682 | UP000007879 |
| *Geodia barretti* (Barrett's horny sponge) | Porifera | 519541 | UP001174909 |
| *Trichoplax adhaerens* | Placozoa | 10228 | UP000009022 |
| *Nematostella vectensis* (Starlet sea anemone) (CH2 X CH6) | Cnidaria | 45351 | UP000001593 |
| *Hydra vulgaris* (Hydra) (Hydra attenuata) (105) | Cnidaria | 6087 | UP000694840 |
| *Paramuricea clavata* (Red gorgonian) (Violescent sea-whip) (A484AB) | Cnidaria | 317549 | UP001152795 |
| *Drosophila melanogaster* (Fruit fly) (Berkeley) | Bilateria | 7227 | UP000000803 |
| *Danaus plexippus plexippus* | Bilateria | 278856 | UP000007151 |

| **Name** | **Group** | **Taxon ID** | **Reference proteome** |
|---|---|---|---|
| *Homo sapiens* | Bilateria | 9606 | UP000005640 |
| *Mus musculus* | Bilateria | 10090 | UP000000589 |
| *Arabidopsis thaliana* | Embryophyta | 3702 | UP000006548 |
| *Nicotiana tabacum* | Embryophyta | 4097 | UP000790787 |
| *Oryza sativa* | Embryophyta | 39947 | UP000059680 |
| *Marchantia polymorpha* | Embryophyta | 3197 | UP000244005 |
| *Brachypodium distachyon* | Embryophyta | 15368 | UP000008810 |
| *Chlamydomonas reinhardtii* | Chlorophyta | 3055 | UP000006906 |
| *Escherichia coli* K-12 | Pseudomonadota | 83333 | UP000000625 |
| *Bacillus subtilis* 168 | Bacillota | 224308 | UP000001570 |
| *Caulobacter vibrioides* NA1000 (C. crescentus NA1000) | Pseudomonadota | 565050 | UP000001364 |
| *Vibrio cholerae* O1 biovar El Tor N16961 | Pseudomonadota | 243277 | UP000000584 |
| *Pseudomonas aeruginosa* UCBPP-PA14 | Pseudomonadota | 208963 | UP000000653 |
| *Porphyridium purpureum* | Rhodophyta | 35688 | UP000324585 |
| *Porphyra umbilicalis* | Rhodophyta | 2786 | UP000218209 |
| *Chondrus crispus* | Rhodophyta | 2769 | UP000012073 |
| *Chlorella variabilis* | Chlorophyta | 554065 | UP000008141 |
| *Ostreococcus tauri* | Chlorophyta | 70448 | UP000009170 |
| *Klebsormidium nitens* | Streptophyta | 105231 | UP000054558 |
| *Chara braunii* | Streptophyta | 69332 | UP000265515 |

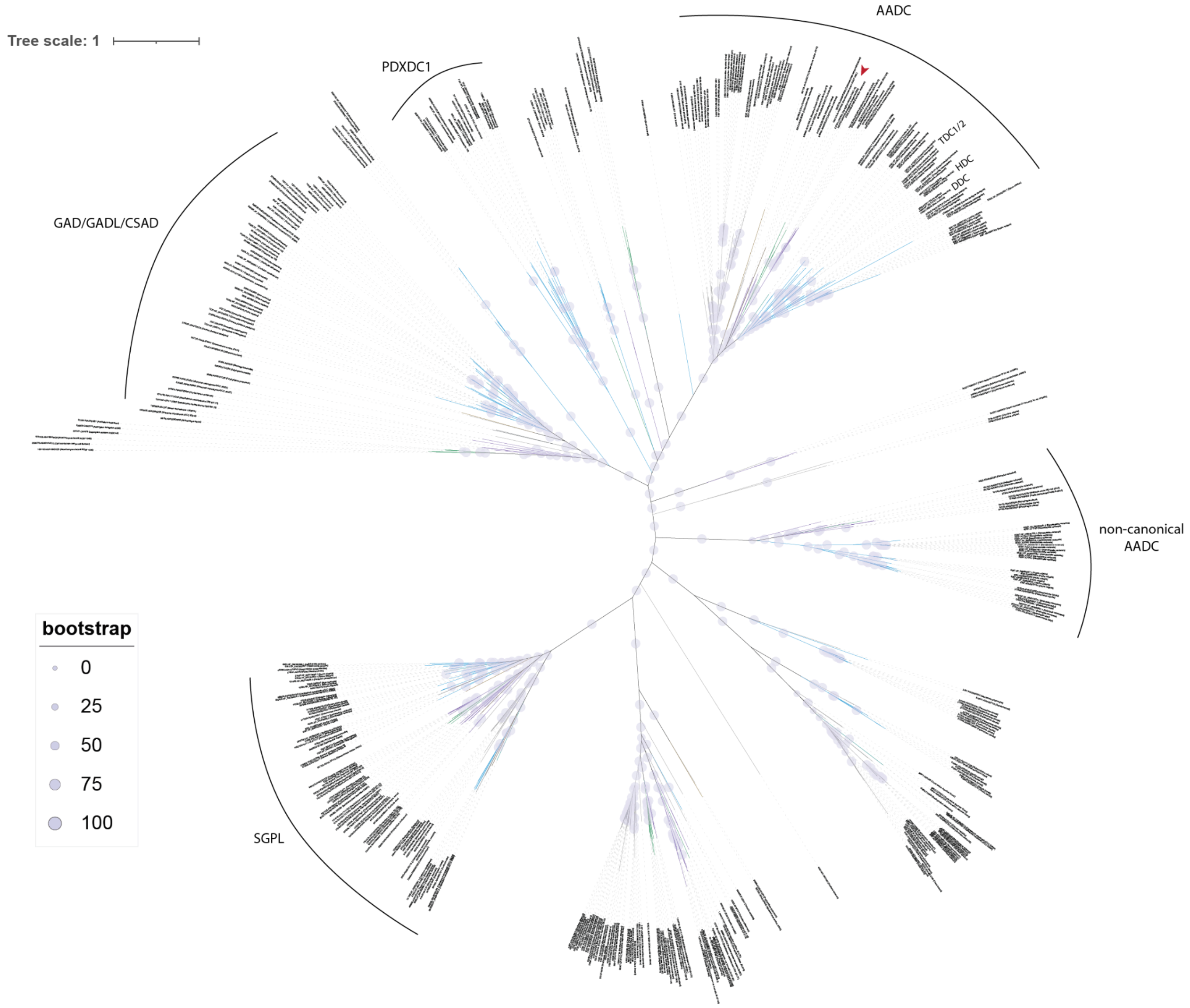


**Fig. S2. Phylogeny of Pyridoxal_deC (PF00282) domain proteins (AADCs).** *Fungi*: purple; *Ascomycota* (within *Fungi*): green; *Metazoa*: blue; *Filasterea*: teal; *Ichthyosporea*: olive brown; *Choanoflagellata*: dark grey; *Fonticula alba*: dark brown; Others: light grey. The cloned fungal sequence is denoted with a red arrow.

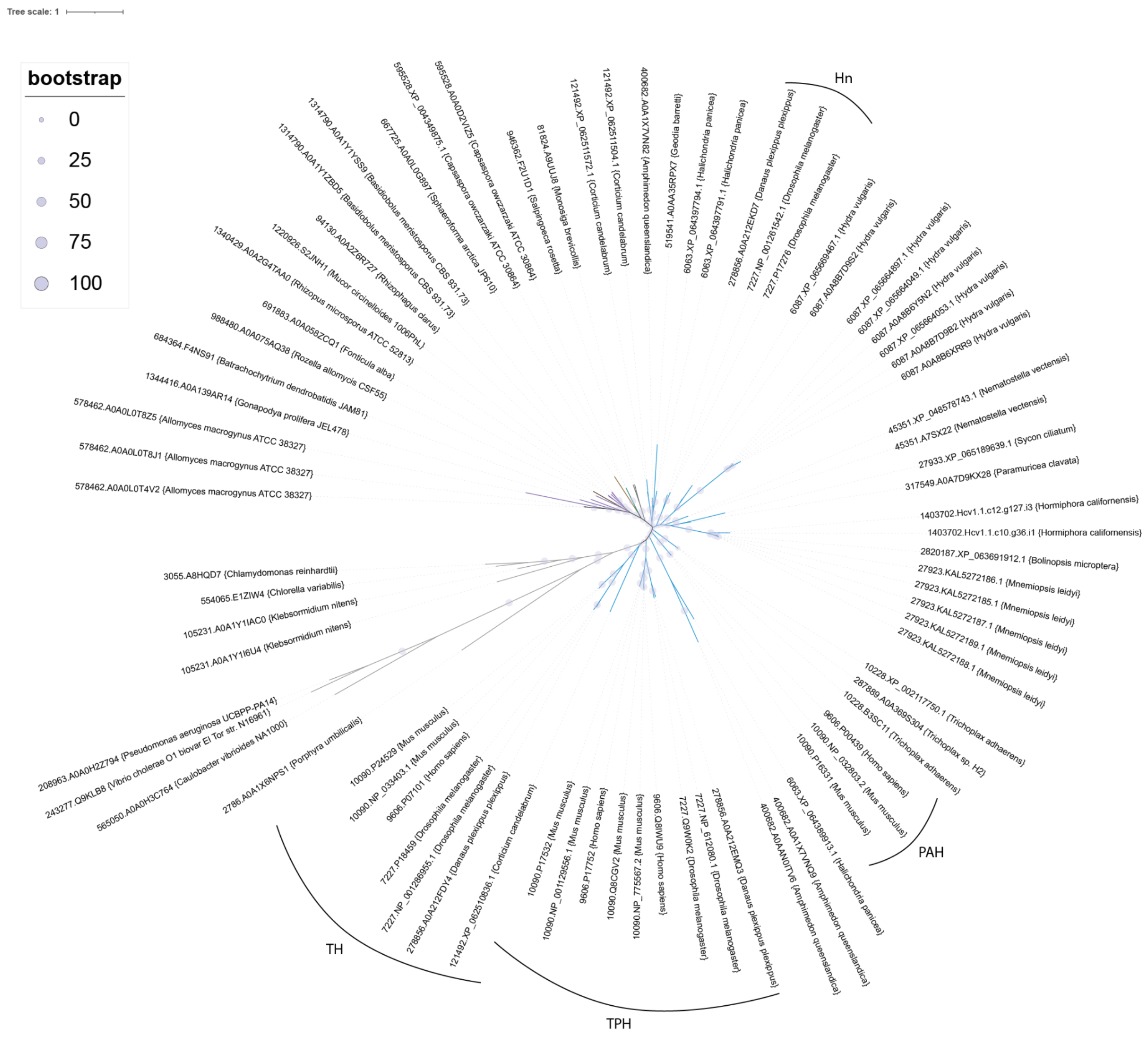

**Fig. S3. Phylogeny of Biopterin_H (PF00351) domain proteins (AAAHs).** Colors are as in fig. S2.

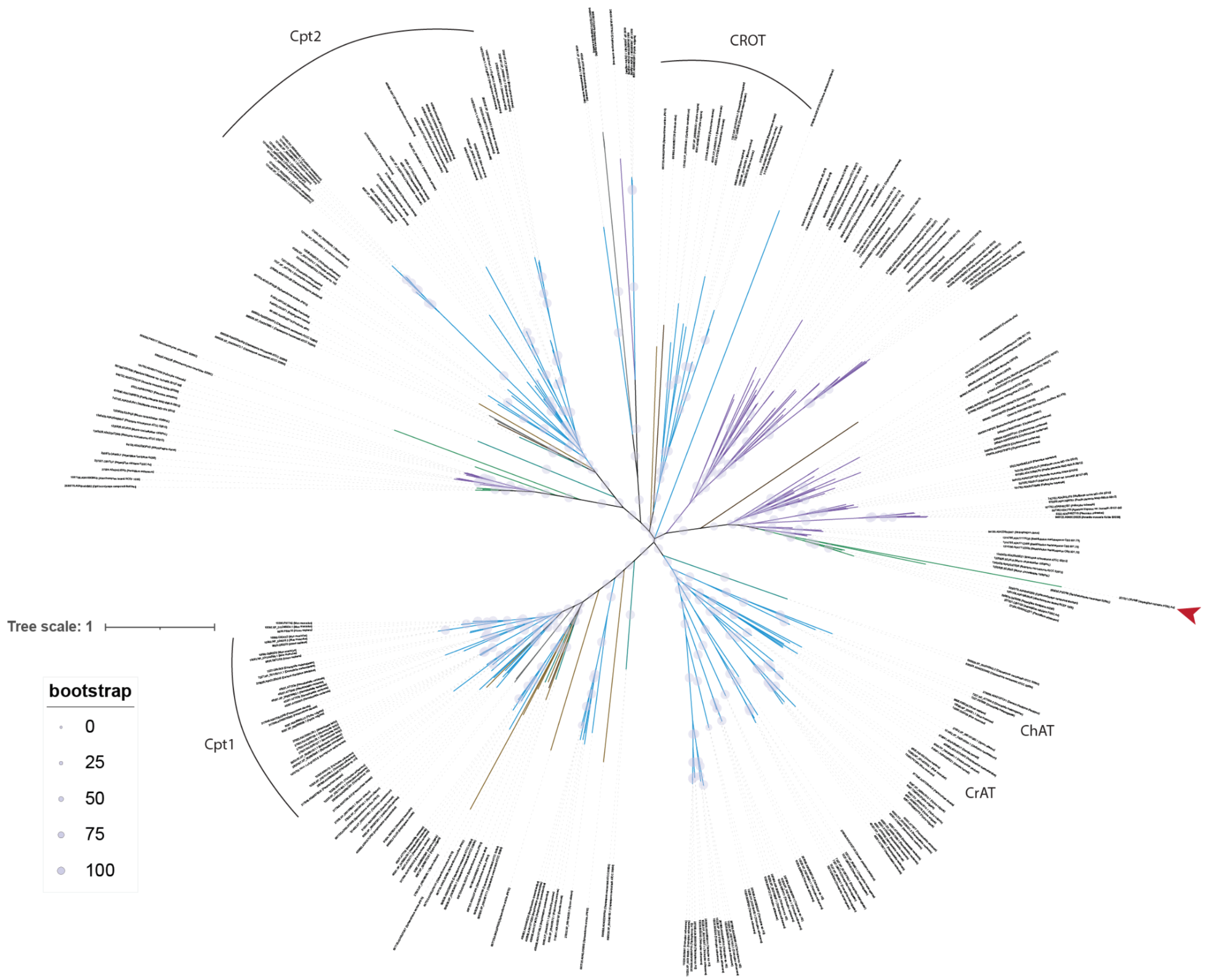


**Fig. S4. Phylogeny of Carn_acyltransf (PF00755) domain proteins (ChAT/CrAT).** Colors as in fig. S2. The cloned fungal sequence is denoted with a red arrow.

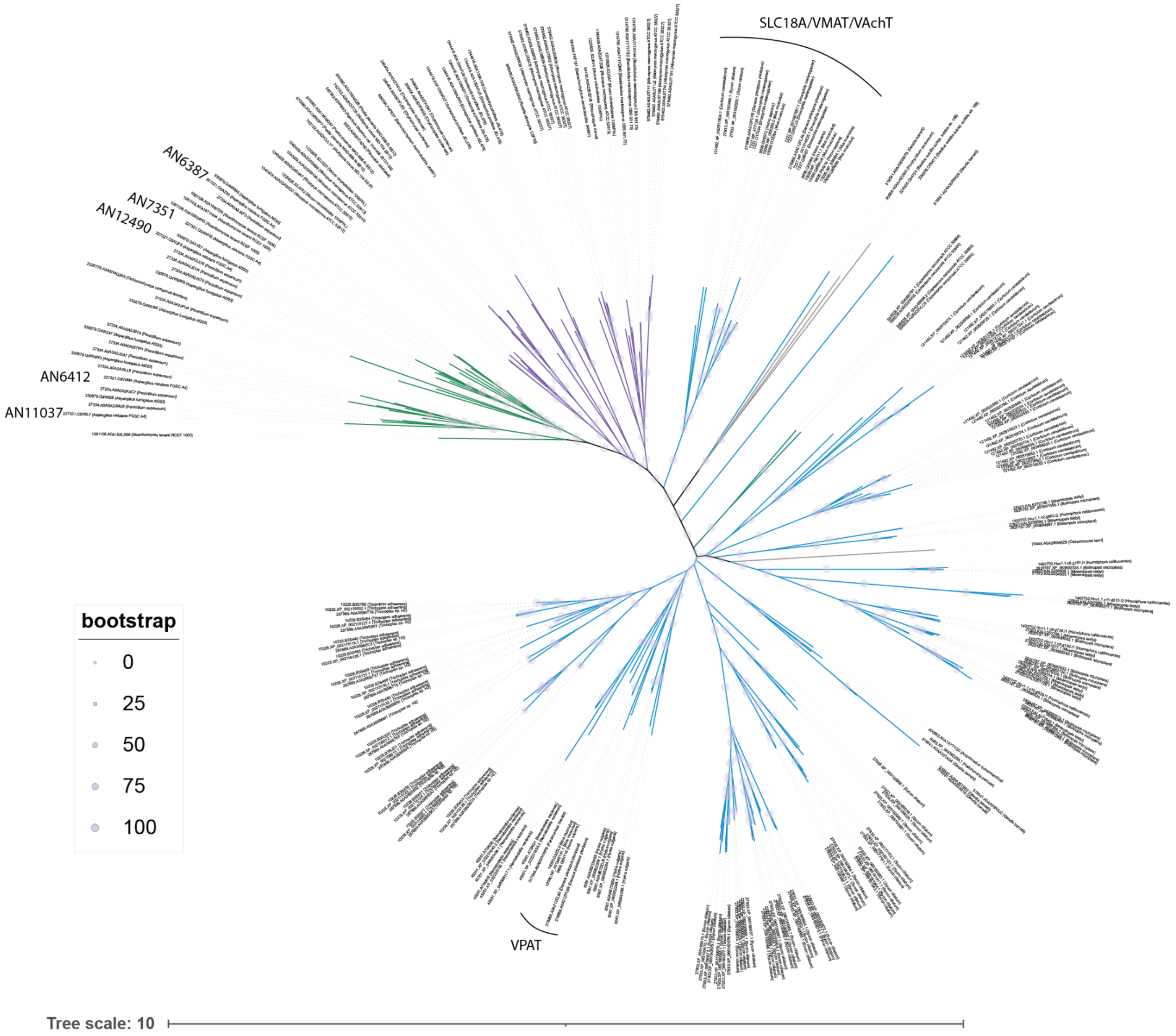


**Fig. S5. Phylogeny of the SLC18 transporter clusters.** Colors as in fig. S2. The gene codes of the cloned transporters are shown next to their leaves.

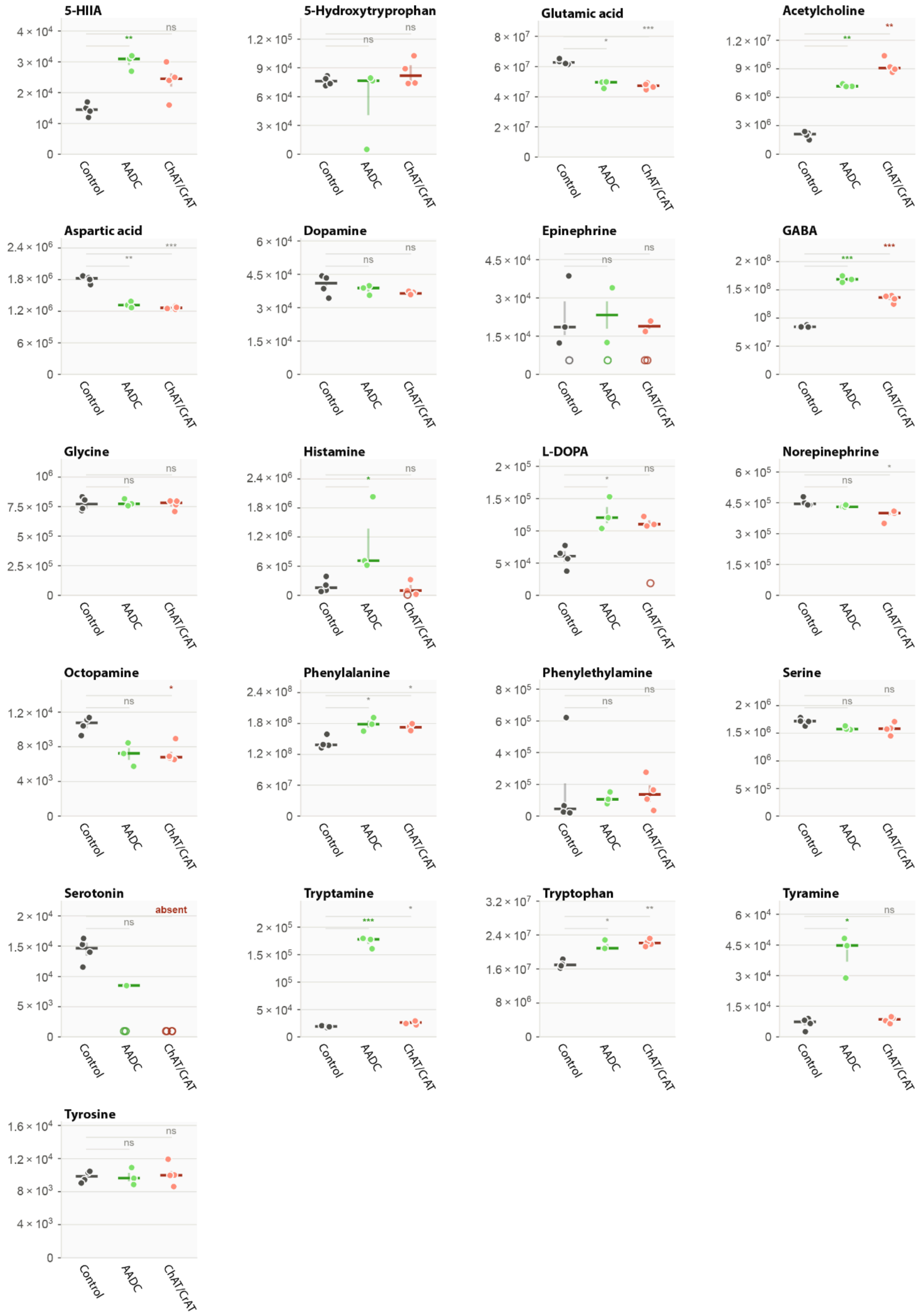
5-HIIA
5-Hydroxytryprophan
Glutamic acid
Acetylcholine
Aspartic acid
Dopamine
Epinephrine
GABA
Glycine
Histamine
L-DOPA
Norepinephrine
Octopamine
Phenylalanine
Phenylethylamine
Serine
Serotonin
absent
Tryptamine
Tryptophan
Tyramine
Tyrosine
Control
AADC
ChAT/CrAT

**Fig. S6. Relative metabolomic quantification of all targeted compounds.** Extended version of Fig. 2C including all compounds measured. The y-axis is the relative peak area. Strains expressing the specified proteins are compared to a control strain. Hollow points: compound not detected in the sample.  Stars = BH q of the Welch test on $log_{10}$ area versus the strain's own control (*** $q < 0.001$, ** $q < 0.01$, * $q < 0.05$); coloured and bold = also at least 1.5-fold and above three times the QC coefficient of variation; grey = significant but below the effect-size or noise threshold.

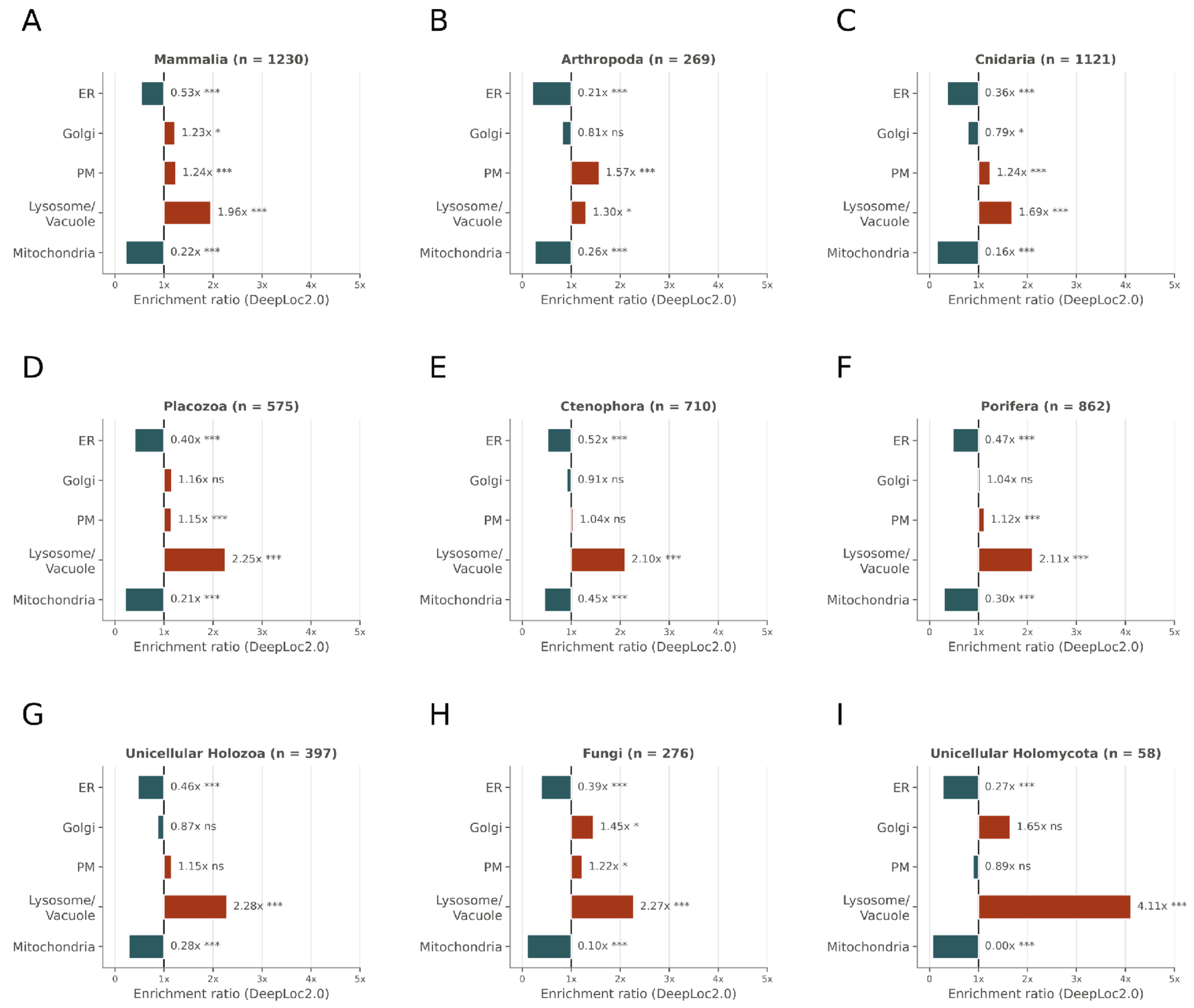


**Fig. S7. Predicted localisation enrichment of transmembrane SV protein homologs across Opisthokonta. A to I.** Ratio of observed localisations to expected localisation for specific organelles for each taxonomic group. Extended version of Fig. 4B, C; n: number of transmembrane SV homologs. Stars denote adjusted q < 0.05 (*), < 0.01 (**) and < 0.001 (***); ns: not significant.

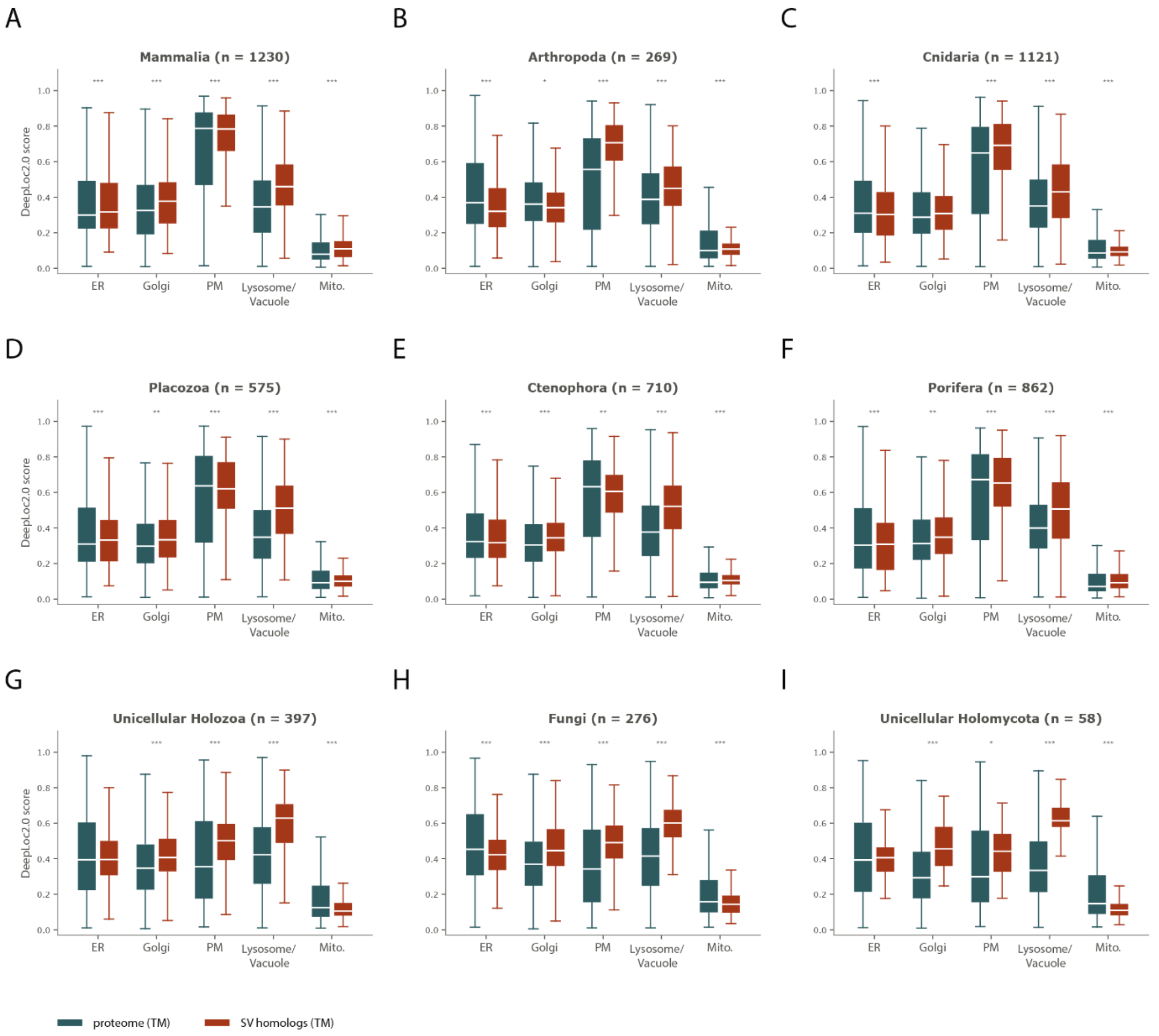


**Fig. S8. Localisation probability distributions of transmembrane SV homologs versus whole transmembrane proteomes across Opisthokonta. A to I.** DeepLoc2.0 localisation probabilities of SV homologs versus whole transmembrane proteomes for each taxonomic group. Extended version of Fig. 4D, E. Stars denote adjusted q < 0.05 (*), < 0.01 (**) and < 0.001 (***).

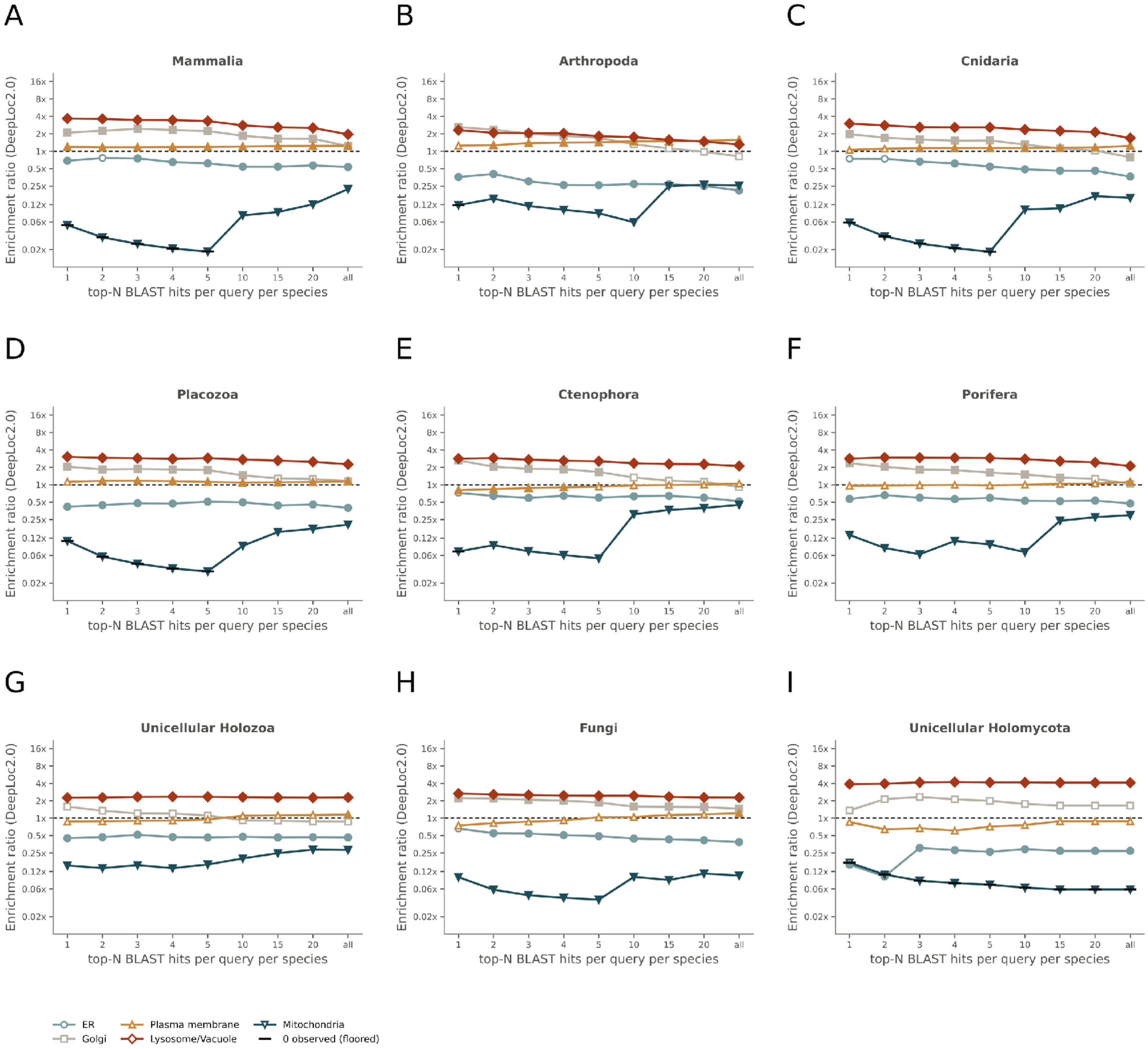


**Fig. S9. Localisation enrichment of transmembrane SV homologs for different numbers of top BLASTP hits per query. A to I.** Ratio of observed localisations to expected localisation for specific organelles for each taxonomic group in a sweep including different numbers of top hits per query. Extended version of fig. 4F, G. Empty plot points: no significant enrichment (methods).

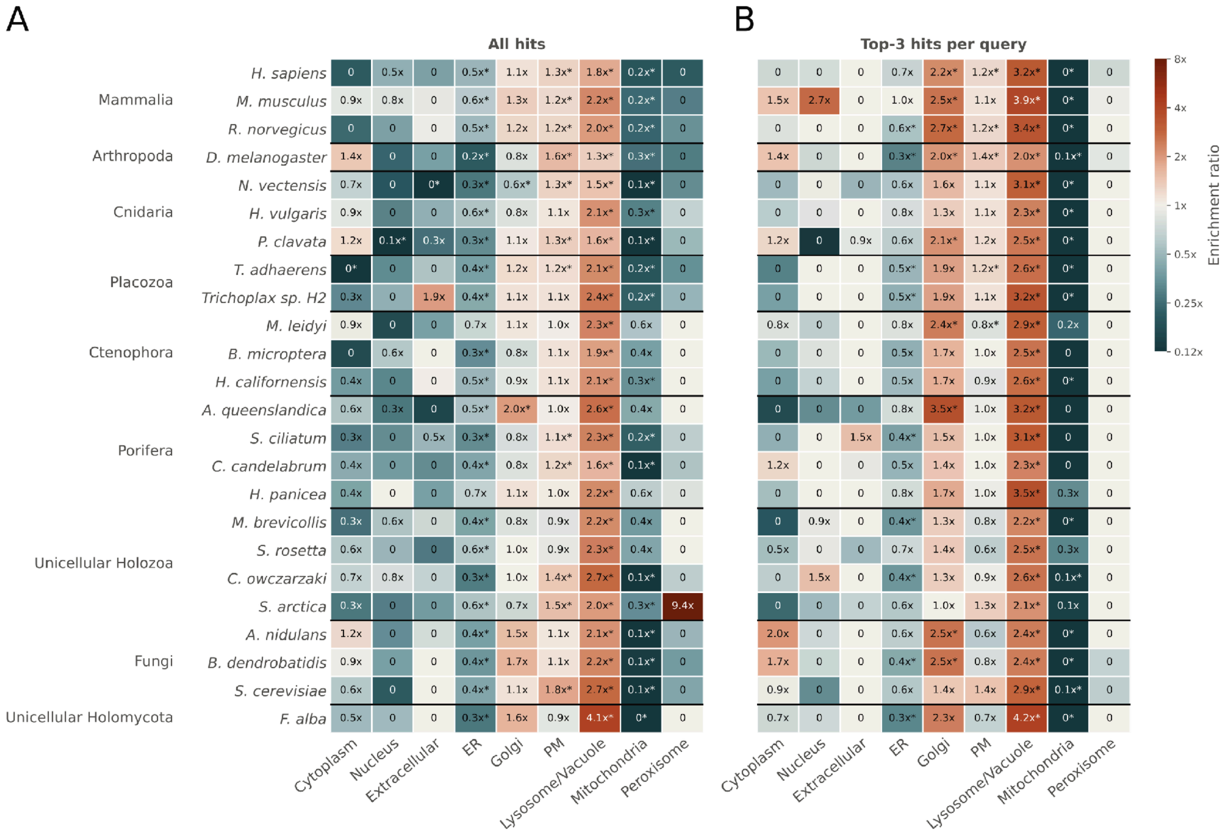


**Fig. S10. Localisation enrichment of transmembrane SV homologs for each species separately. A.** All BLASTP hits per query. **B.** Top-3 hits per query. Colour denotes the ratio of observed localisations to expected localisation. Asterisks denote adjusted $q < 0.05$.

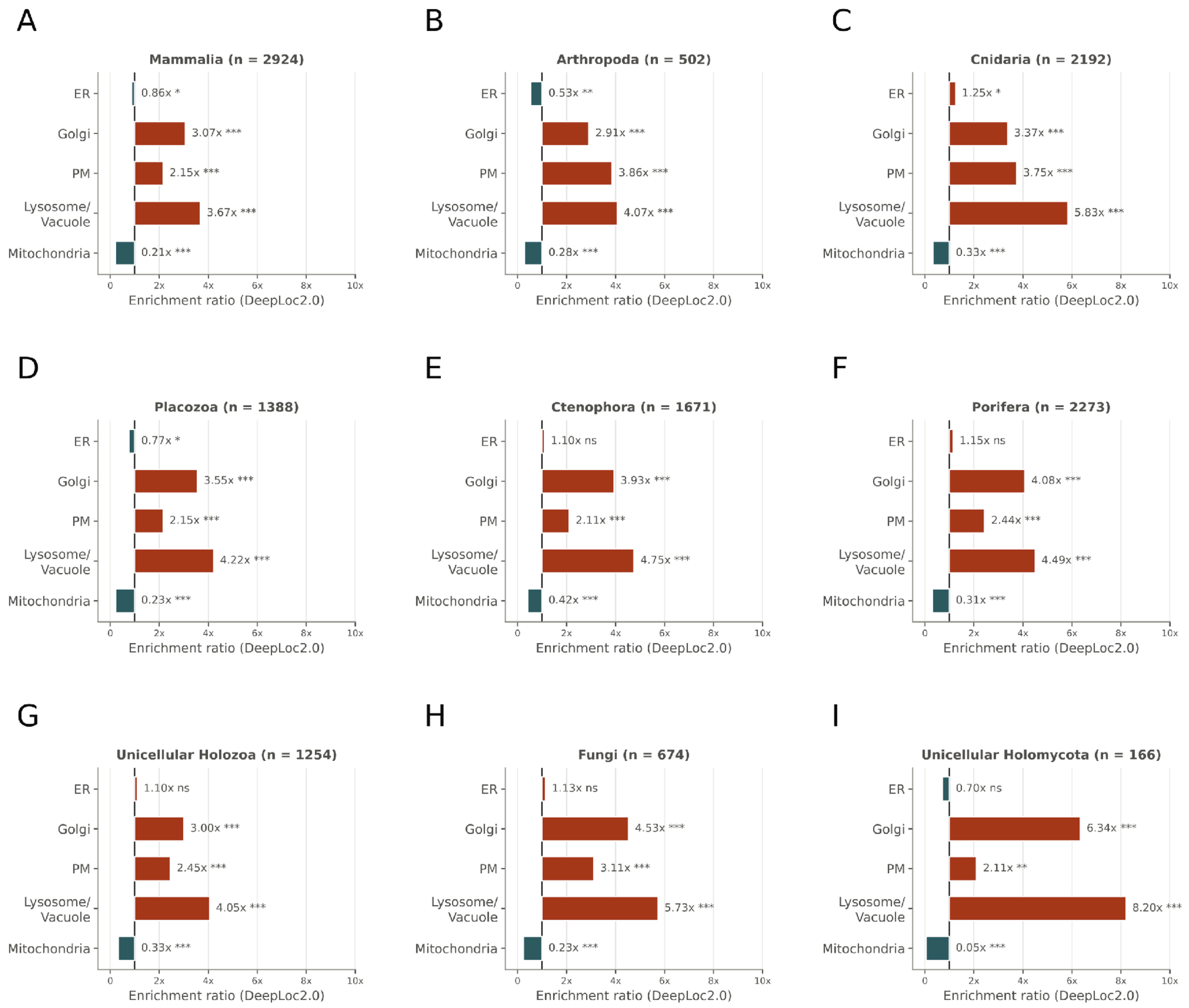


**Fig. S11. Predicted localisation enrichment of SV protein homologs across Opisthokonta without transmembrane filtering. A to I.** Same as fig. S7, but all SV homologs are compared to whole proteomes. Stars denote adjusted q < 0.05 (*), < 0.01 (**) and < 0.001 (***); ns: not significant.

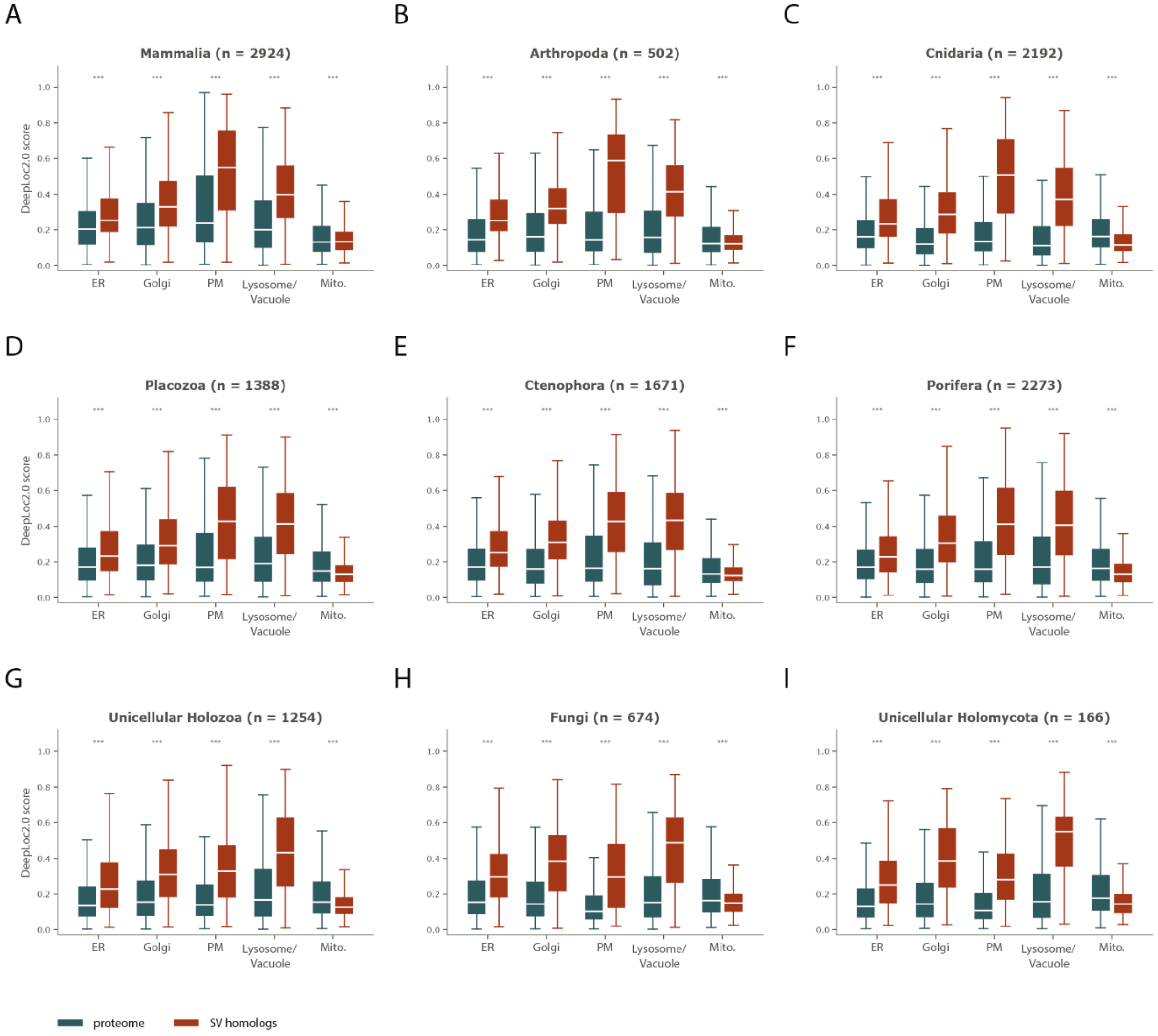


**Fig. S12. Localisation probability distributions of SV homologs versus whole proteomes without transmembrane filtering. A to I.** Same as fig. S8, but all SV homologs are compared to whole proteomes. Stars denote adjusted q < 0.05 (*), < 0.01 (**) and < 0.001 (***).

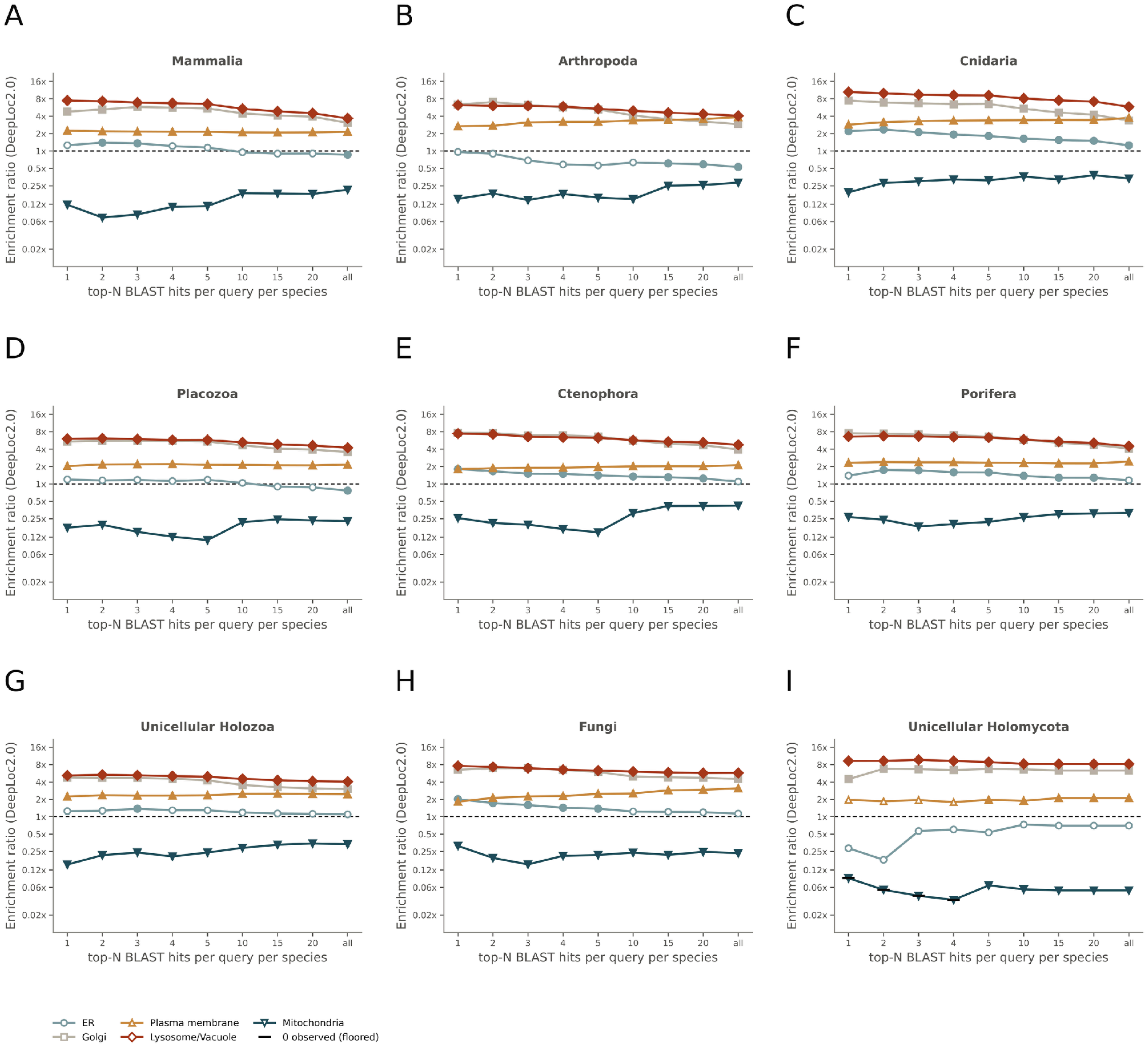


**Fig. S13. Localisation enrichment of SV homologs for different numbers of top BLASTP hits per query without transmembrane filtering. A to I.** Same as fig. S9, but all SV homologs are compared to whole proteomes. Empty plot points: no significant enrichment (methods).

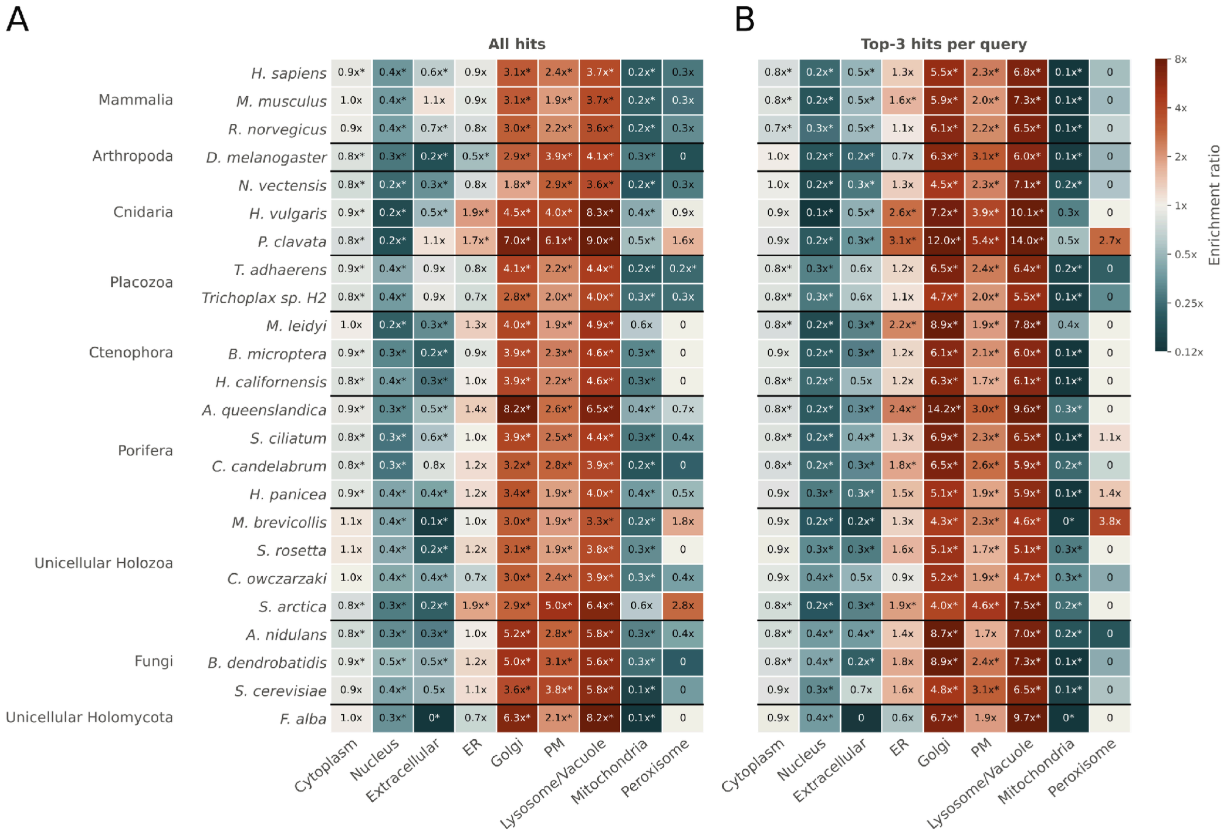


**Fig. S14. Localisation enrichment of SV homologs for each species separately without transmembrane filtering. A.** All BLASTP hits per query. **B.** Top-3 hits per query. Same as fig. S10, but all SV homologs are compared to whole proteomes. Asterisks denote adjusted q < 0.05.

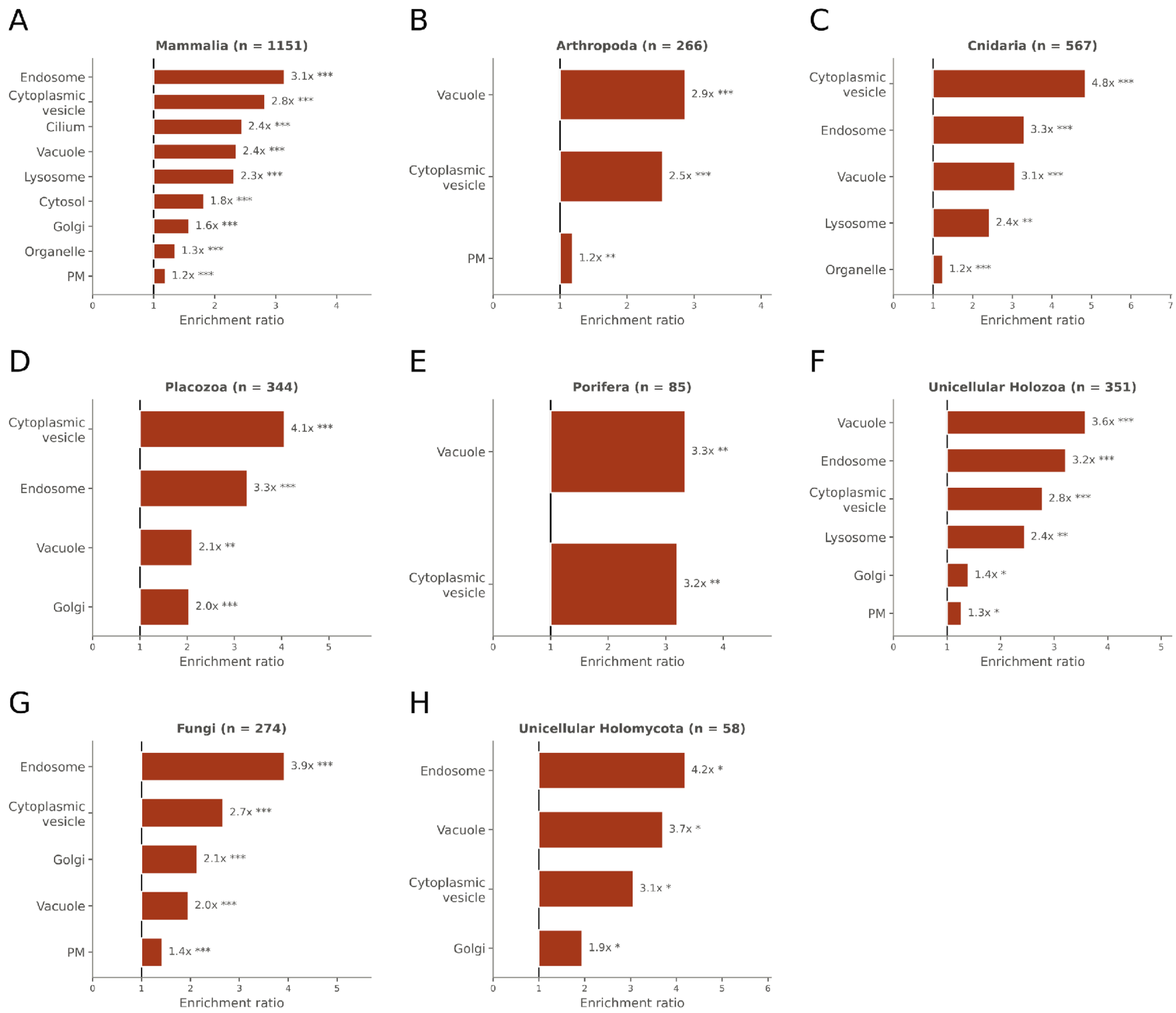


**Fig. S15. GO slim (cellular component) enrichment for transmembrane SV homologs across Opisthokonta. A to H.** Enrichment of SV homolog term counts versus expected term counts for each taxonomic group. Only proteins with GO annotation were used; Ctenophora are absent because no GO annotation was available. Extended version of fig. 4H, I. Stars denote adjusted q < 0.05 (*), < 0.01 (**) and < 0.001 (***).

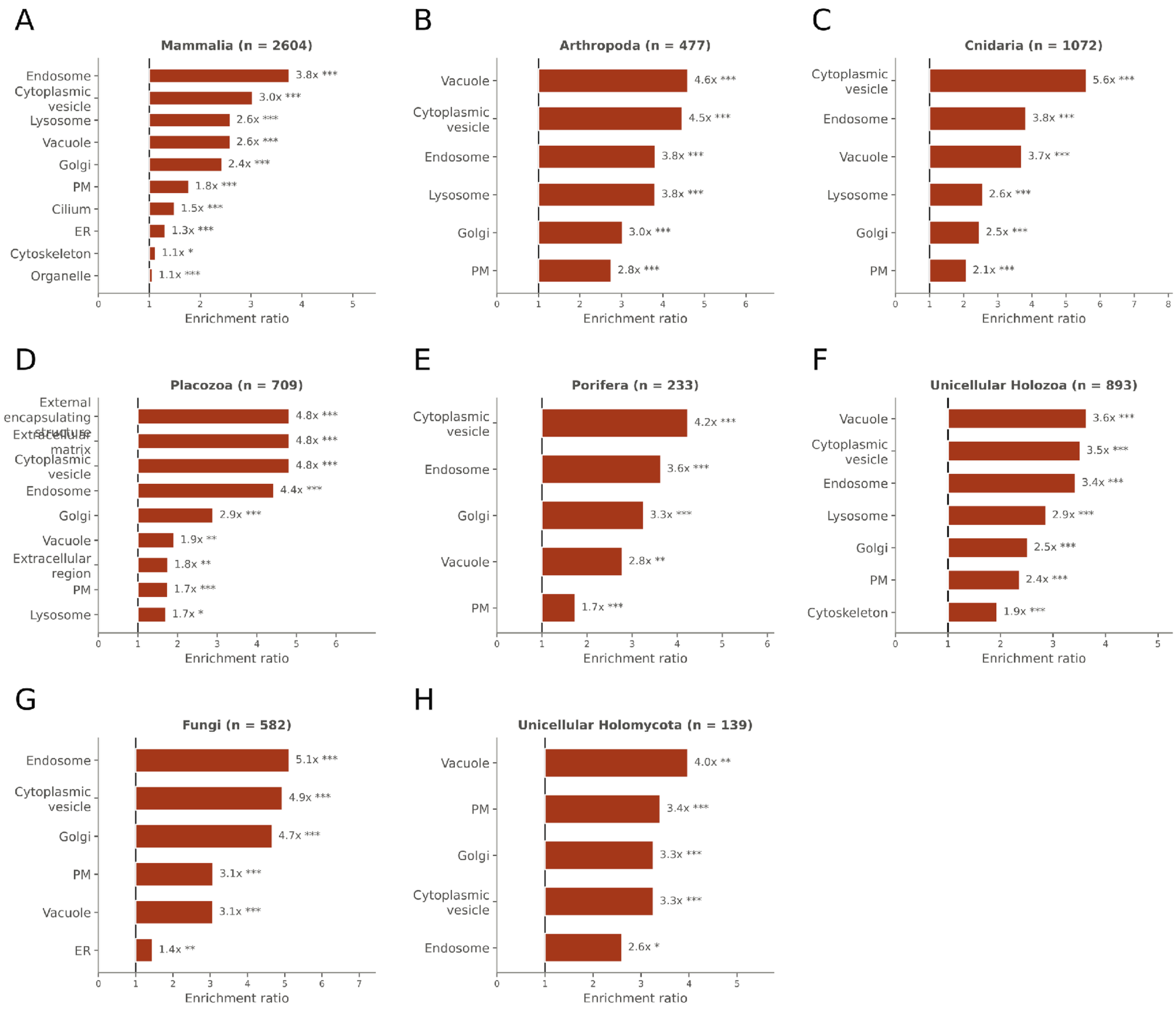


**Fig. S16. GO slim (cellular component) enrichment for SV homologs without transmembrane filtering. A to H.** Same as fig. S15, but all SV homologs are compared to whole annotated proteomes. Stars denote adjusted q < 0.05 (*), < 0.01 (**) and < 0.001 (***).

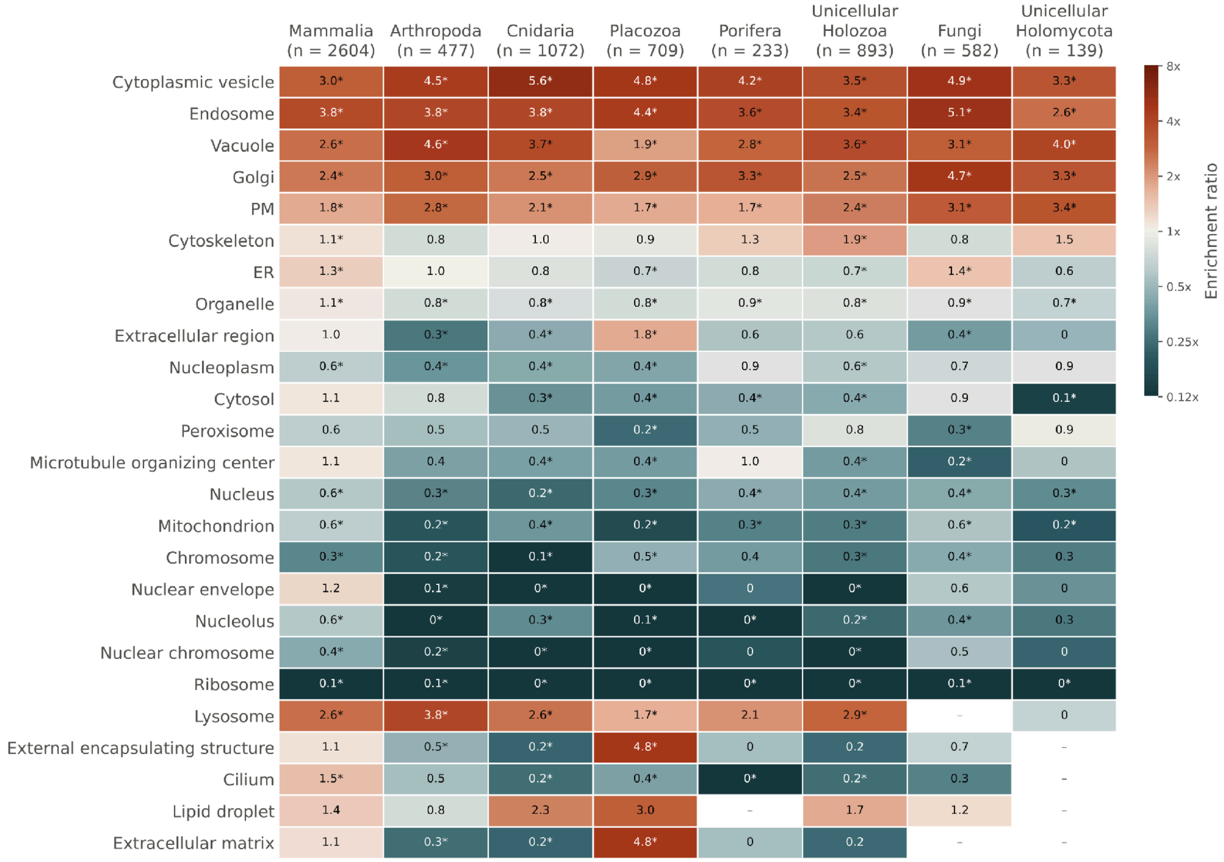


**Fig. S17. GO slim (cellular component) enrichment ratios for all tested terms without transmembrane filtering.** Heatmap of SV homolog term counts versus expected term counts for each taxonomic group; n: number of annotated SV homologs. Asterisks denote adjusted q < 0.05; dashes: term not tested.

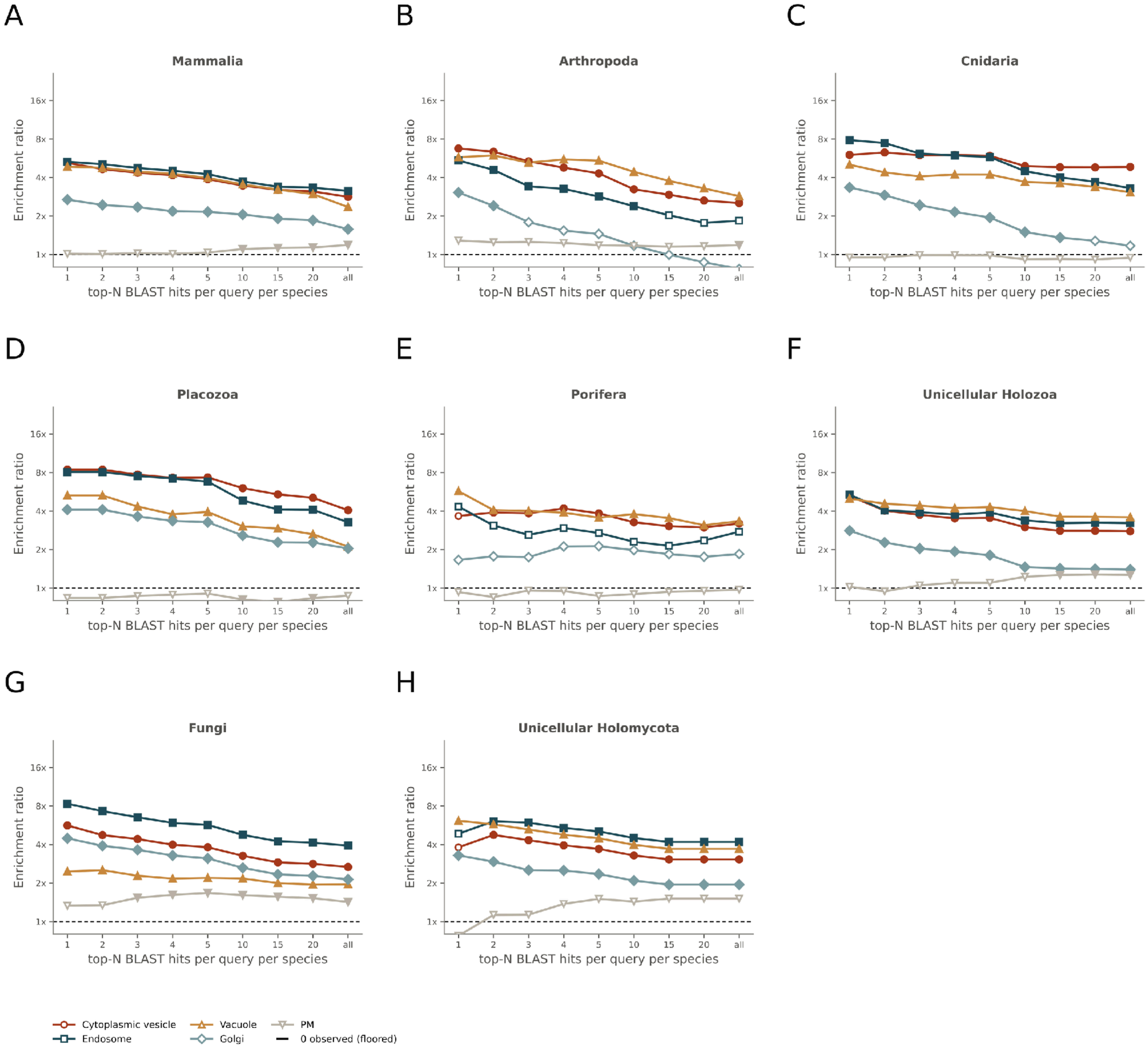


**Fig. S18. GO slim (cellular component) enrichment for different numbers of top BLASTP hits per query. A to H.** Enrichment ratio of the top enriched terms for each taxonomic group in a sweep including different numbers of top hits. Empty plot points: no significant enrichment.

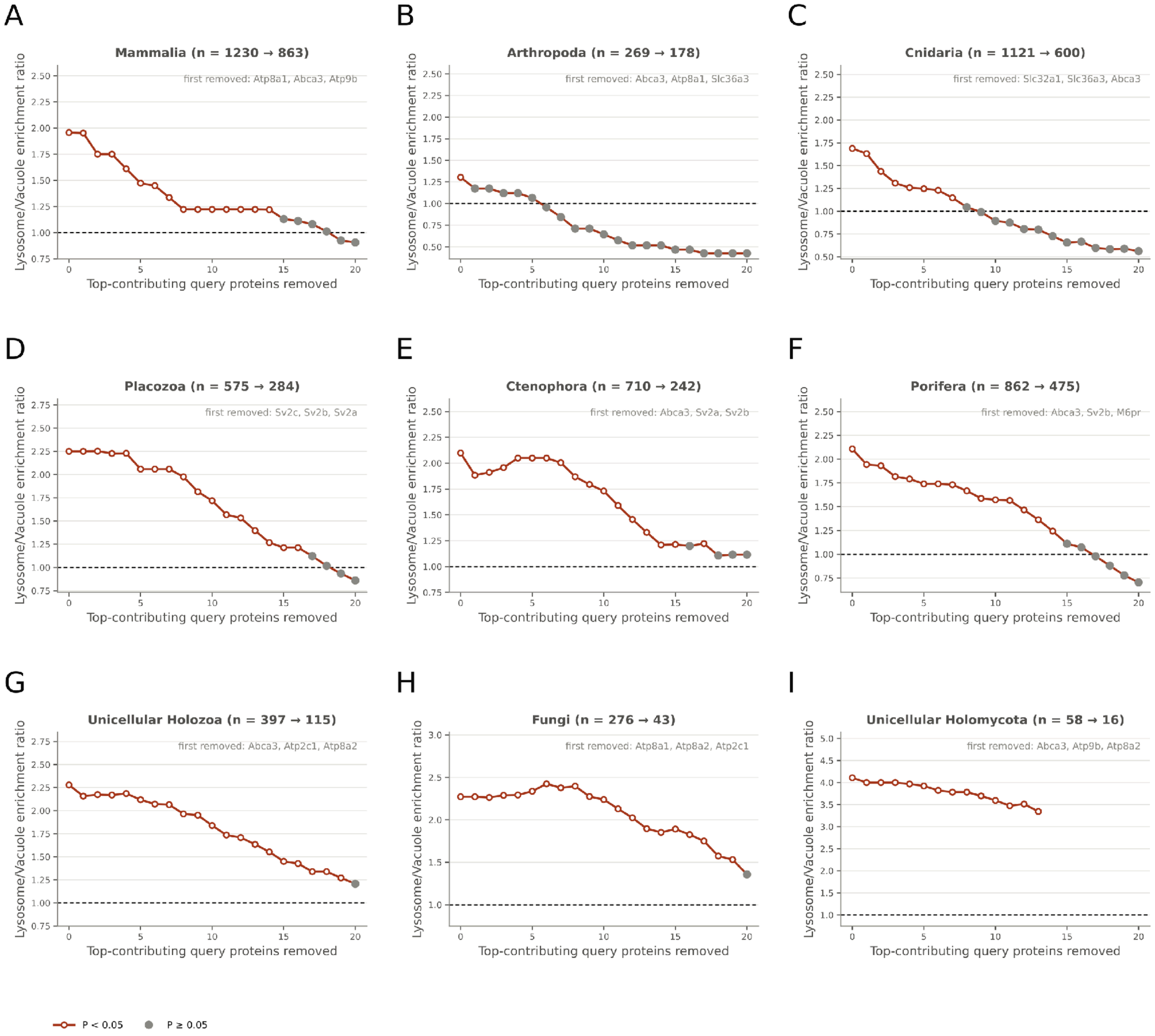


**Fig. S19. Query jackknife of the lysosome/vacuole enrichment. A to I.** Lysosome/vacuole enrichment ratio of transmembrane SV homologs after cumulative removal of the query proteins contributing the most lysosome/vacuole-predicted homologs, for each taxonomic group. n: number of homologs before and after removal of 20 queries; the first three queries removed are noted. Red points: $P < 0.05$; grey points: $P \geq 0.05$.

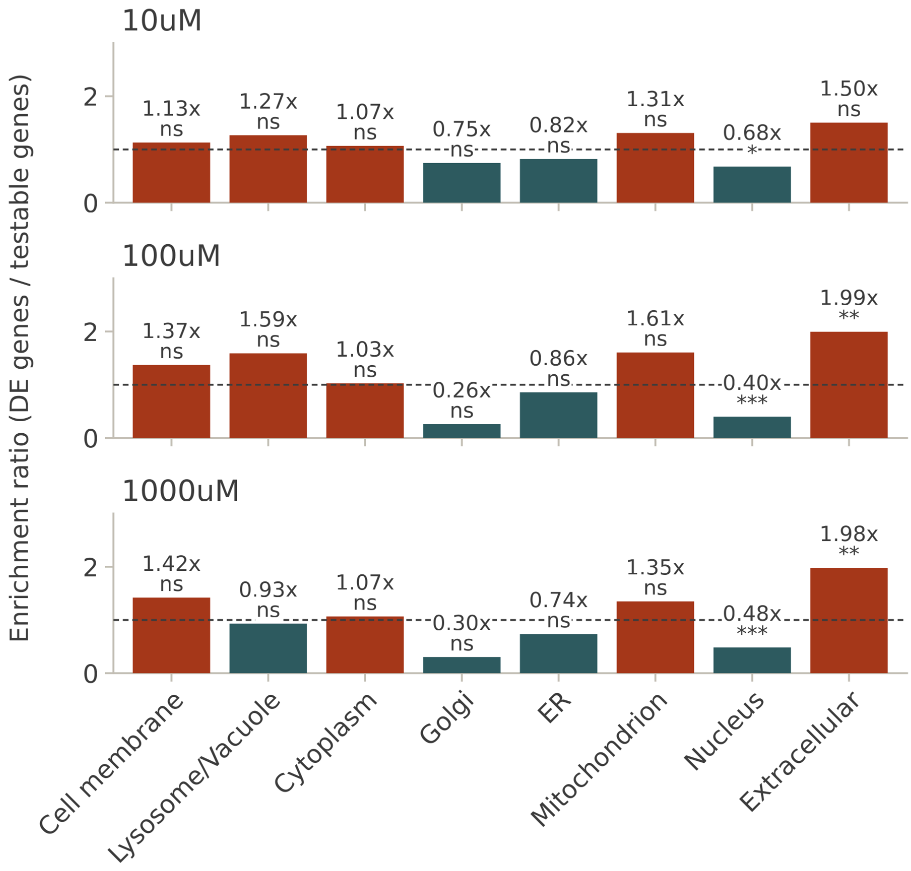


**Fig. S20. Downregulated DE genes are enriched for extracellular proteins.** Enrichment of downregulated DE genes versus the testable proteome for predicted localisation ratios of their proteins, for each trace amine concentration. Same as fig. 5E for downregulated genes. Stars denote adjusted q < 0.05 (*), < 0.01 (**) and < 0.001 (***); ns: not significant.

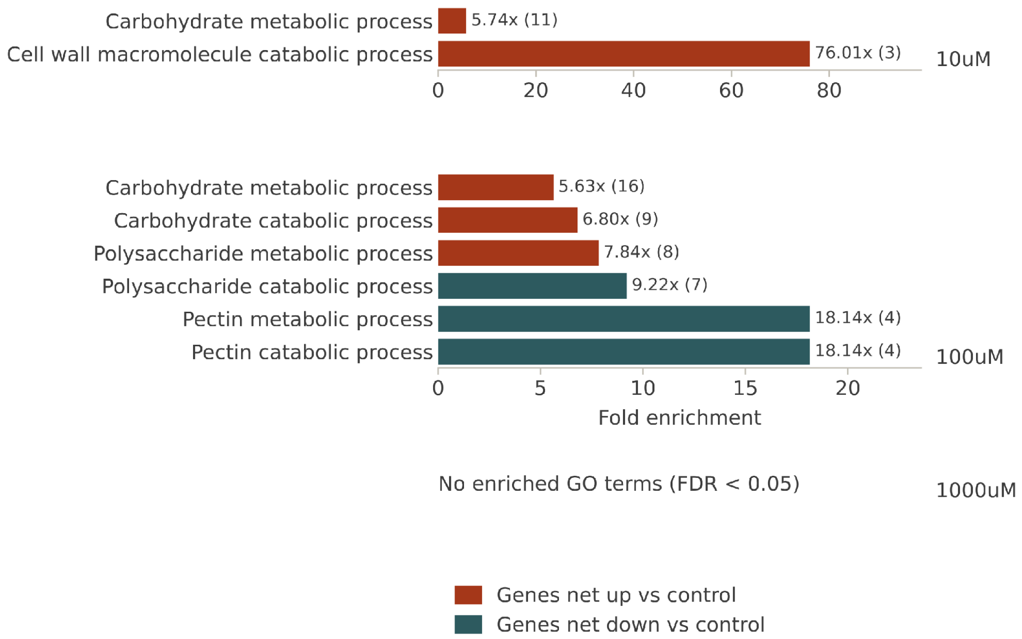


**Fig. S21. GO enrichment (biological process) of extracellular DE genes.** Fold enrichment of GO terms for DE genes (up- and downregulated together) predicted to be extracellular, versus the testable genes, for each trace amine concentration. The number of DE genes per term is in parentheses. Colour denotes whether the genes of each term are net up- or downregulated versus the control. No terms were enriched at 1000 μM.
